# Similar structural area modulates local inhibition initiating post-lesion adaptive mechanism

**DOI:** 10.1101/2023.03.07.531541

**Authors:** Priyanka Chakraborty, Suman Saha, Gustavo Deco, Arpan Banerjee, Dipanjan Roy

## Abstract

The focal lesion, a form of biological perturbation damaging anatomical architecture, reasonably alters the normative healthy functional pattern but may recover over time. Nevertheless, how the brain counters deterioration in structure by global reshaping of functional connectivity (FC) after a lesion is largely unknown. We propose a novel equivalence principle based on structural and dynamic similarity analysis to predict specific compensatory areas initiating lost excitatory-inhibitory (E-I) regulation after lesion. We hypothesize that similar structural areas (SSAs) and dynamically similar areas (DSAs) corresponding to a lesioned site are the crucial dynamical units to restore lost homeostatic balance within the surviving cortical brain regions. SSAs and DSAs are independent measures, one based on structural similarity properties measured by Jaccard Index and the other based on post-lesion recovery time. Thereafter, a large-scale mean field model is deployed on top of a virtually lesioned structural connectome for characterizing the global brain dynamics and functional connectivity at the level of individual subjects. Despite inter-individual variability in SSAs, we found a general normative pattern in functional re-organization within the ipsi- and contra-lesional regions. The study demonstrates how SSAs and DSAs largely predict overlapping brain regions for different lesion centers/sites irrespective of the complexity of the lesion recovery process. The proposed computational framework captures the improvement of large-scale cortical cohesion by re-adjusting local inhibition. Our results further suggest that the predicted brain areas participating in recovery are not randomly distributed and widespread over the brain. Instead, the predicted brain areas are predominantly recruited from the ipsilesional hemisphere, barring a few regions from contra, suggesting that wiring proximity and similarity are the two major guiding principles of compensation-related utilization of hemisphere (CRUH) in the post-lesion FC re-organization process. Our finding further suggests that the re-organization of FC arises from the interplay between the underlying structural connectivity profile and the local inhibitory weights influencing compensatory coordinated brain dynamics during post-lesion recovery.

## I. INTRODUCTION

One of the fundamental queries in neuroscience is ‘How does the brain adapt to post-lesion recovery and whether some general normative patterns exist irrespective of individual variations in lesion extent and locations? What are compensatory mechanisms critical to brain network recovery during post-injury recovery, including changes in local homeostasis and widespread coordinated cortical activity? Here, we propose a detailed computational framework using structural-and-functional equivalence principles to demonstrate that the brain’s normative spontaneous dynamical pattern is compensated by restoring local homeostasis post-lesion and whether the participating compensatory areas are primarily recruited utilizing two major guiding principles, structural similarity and wiring proximity of compensation-related utilization of hemisphere (CRUH) in the post-lesion functional re-organization.

The term ‘focal lesion’ [1, 2] refers to biological perturbation to the anatomical architecture, e.g., damage of a region due to stroke (ischemic stroke due to atherosclerosis, hemorrhagic stroke) [2], traumatic brain injury (TBI) [3], glioma [4] can qualitatively alter short- and long-term brain functions. In lesions, neurons that are deprived due to lack of oxygen, and energy from standard metabolic substrates, cease to function in seconds and show severe signs of anatomical damage after 2 minutes [5]. In the first few days or weeks after injury, regular patterns of synaptic activity in peri-infarct [6–9] and even distant functionality-related structure are interrupted [10]. Failures in the energy-dependent processes due to loss of inputs from adjacent tissue [11] lead towards cell death [12], abnormal neuronal firing rates [13], and may even lead to delayed neuronal injury [14] which inflict local to global level excitation-inhibition (E-I) balance on the neuronal network [15, 16]. These mechanical and cellular alterations can cause chronic functional disabilities, including motor deficits (e.g., hemiparesis), sensory (e.g., hemianopia), and higher-order cognitive processes (e.g., aphasia, hemispatial neglect) [17] and abnormal movement synergies [18, 19].

Studies have revealed the mechanism for lesion recovery and identified associated factors in primate and nonprimate [20, 21]. For example, the cerebral cortex triggers a plastic mechanism in adjacent and remote areas in post-lesion phases, correlated with limited, spontaneous restoration of function [22–24]. Two significant factors are involved in the plasticity mechanism in lesion recovery [25], (i) an amount of surviving diffuse and redundant connectivity in the central nervous system, and (ii) new functional circuits can form through remapping between related cortical regions. Homeostatic plasticity is a negative feedback-mediated form of plasticity, also known as synaptic scaling [25], that keeps network activity at the desired set point [26]. It helps maintain a stable ratio of excitation and inhibition and sustains the desired working point. Nevertheless, local E-I homeostasis engenders functional recovery by increasing excitation and attenuating inhibition in both perilesional and distant cortical areas [10, 27, 28]. In addition, enhancement of cortical excitability in surviving cortical areas would compensate for the lost structural circuits [29] and functional deficits [30–32].

Other key investigations suggest that graph theoretical properties of structural and functional networks plays a crucial role in capturing several aspects of lesion-induced alterations in topological properties of largescale structural and functional brain networks [33–35]. However, these studies have yet to systematically investigate region-specific roles in post-lesion functional restoration of brain networks. A recent longitudinal study on mTBI showed notable changes in structural and functional brain networks in the post-lesion recovery phase [36]. However, they did not identify specific regions participating in the functional recovery process. They found no association between damaged functional and structural connections after TBI [36]. Previous studies have reported network properties, such as nodal strength, participation coefficient, and modularity, played a decisive role in finding the short-term and long-term effects of lesion [37, 38]. However, the region-specific role of anatomical networks in association with the post-lesion global functional recovery still needs to be fully uncovered and remains an open question. From a dynamical systems perspective, the brain is a spatially organized system [39] with time-dependent signal propagation along multiple pathways, each capable of adapting to changes in transmission fidelity [25]. Thus, brain dynamics is governed by underlying anatomy, and the underlying intrinsic biological parameters [40]. However, regional specificity in association with the intrinsic parametric role must be elucidated as to how specific regions may play a vital role by adapting intrinsic parameters in shaping emerged brain dynamics to compensate for structural damage following lesions. With this knowledge gap and motivation, we hypothesize that regions with similar incoming and outgoing connections corresponding to a lesion site, labeled as similar structural areas (SSAs) in this study, could be the potential candidate for re-establishing E-I balance (at the level of both local and global brain scale) in the postinjury period. Hence, the prediction of SSAs is one of the fundamental contributions of this study. The second fundamental contribution to identifying dynamically similar areas (DSAs) using readjustment time, a dynamic measure indicating the re-establishment of local E-I balance post-lesion. For example, a region that helps compensate for motor deficit should get incoming motor information from its adjacent or distant areas, thus would be functionally relevant and structurally equivalent. SSA may form complementary or redundant connections in the surviving areas, providing an alternative pathway for information fidelity after the lesion. Thus, SSAs could lead the adaptive mechanism to compensate for lost local homeostasis and inter-areal excitability, further reshaping collective activity.

We have systematically addressed the following questions: (i) What changes in E-I balance cause altered neural activity after early brain injury due to anatomical network damage? (ii) Which are the notable areas that re-adjust their inhibitory weights to balance E-I homeostasis and sustain a target firing rate *∼*4Hz? (iii) What processes are related to the post-injury functional re-organization within the surviving structural network? To address these questions, we Identify the factors, e.g., time to re-establish E-I balance in local areas and modulated local inhibitory weights, next we find signatures from the structural properties in correlation with coordinated neural dynamics, and finally, identify the mechanisms displaying correlations between the parameters controlling local E-I homeostasis and structural network similarity measure, e.g., the Jaccard coefficient. Our approach identifies SSAs and DSAs (two independent measures) predicting essentially similar brain regions that participate in the compensation-related utilization of the hemisphere (CRUH) on the road to recovery. These areas reset local E-I balance post-lesion by modulating their inhibitory weights. Our findings suggest a functional network recovery process could be fully predicted based on SSAs. The high correlation between these two mutually exclusive methods (SSAs and DSAs) arises from the interplay between the underlying structural connectivity profile and the local inhibitory weights influencing compensatory coordinated brain dynamics during post-lesion recovery.

## II. RESULTS

To test our hypothesis, we simulate a virtual lesion model. The virtual lesion is introduced by deleting incoming and outgoing connections of a node in the structural connectome of healthy subjects [33, 37]. Fig. 1a shows a large-scale dynamical mean field (DMF) model [41, 42] on top of the virtually lesioned structural connectivity (SC), labeled as the virtual lesion model. A feedback inhibition control (FIC) algorithm [41], a negative feedback-mediated form of plasticity, re-adjusts local inhibition synaptic weights to restore E-I balance and target firing rate of *∼*4Hz [43] in the post-injury period.

**FIG. 1.**
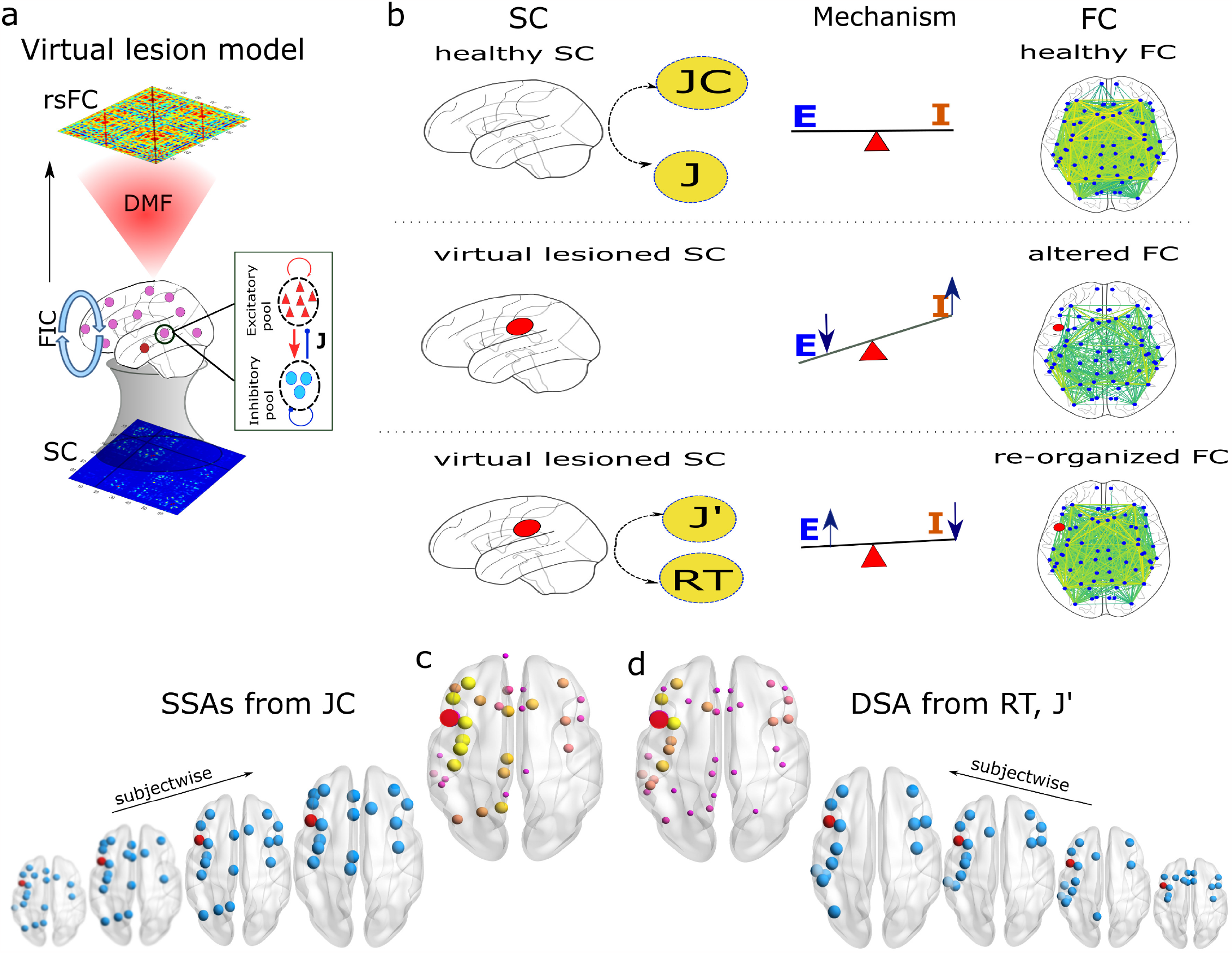
Workflow. (a) Schematic of the virtual lesion model. SC is generated based on the Desikan-Killiany atlas with 68 regions of interest (ROI). A dynamical mean field model (DMF) is spatially connected via virtually lesioned SC. The DMF model generates resting-state neural activity using a feedback inhibition control (FIC) algorithm. Synthetic BOLD series are generated from the neural signals using a hemodynamic model. The FIC is a recursive process of adjusting local inhibitory feedback weight (*J*). At optimal conditions, each region maintains balanced homeostasis and an excitatory firing rate between 2.63-3.55Hz. The pairwise Pearson correlation coefficient determines the simulated resting-state FC. (b) Our analysis is performed considering three conditions, placed in three rows. The top row shows healthy conditions. First, the Jaccard coefficient (*JC*) matrix is calculated from a healthy SC. Next, the DMF model coupled via healthy SC has been run with the FIC algorithm establishing E-I balance. The final value of each brain region’s feedback synaptic inhibitory weights (*J*) is stored beside the synaptic activity. The synaptic gating variables are further processed to generate model-based rsFC. The seesaw represents the status of the global E-I balance state. The obtained *J* have been used further as initial values for inhibitory coupling weights in the model simulation for the rest of the two conditions. In the second row, a virtual lesion is introduced to the SC by setting all rows and columns equal to zero. Next, the virtual lesion model is run without the FIC to generate model-based altered FC. The seesaw represents an imbalanced E-I state as an early impact of the lesion. The lower row depicts the third condition when E-I balance is restored in the brain. The virtual lesion model is simulated with the FIC algorithm. Model parameters, such as modulated local synaptic inhibitory weights (*J’*) and re-adjustment time (*RT*), are stored for further analysis. Yellow circles show the model parameters of our interest. Two representative results are shown in (c) and (d). (c) Identified SSAs are shown for individual subjects and group levels. (d) Estimated DSAs using *RT* and *J* ^*‘*^ are shown.

We did not consider other virtual lesion types, such as edge deletion or multi-region damage [33, 37]. The preprocessed data of 49 healthy subjects are taken from the Berlin data set [44]. Desikan-Killiany [45] parcellation divides the brain into 68 regions of interest.

The mathematical framework is set for three conditions based on the SC status and the E-I balance state. Three rows display the three operant conditions for model simulation in Fig. 1b.

The top row in Fig. 1b describes the healthy condition when SC remains intact, and the E-I balance is maintained. We derive the JC matrix from the healthy SC. The ‘Similar Structural Areas’ (SSAs) correspond to a given lesion site from a healthy individual’s SC based on the JC measure before lesion occurrence. We store the local inhibitory weights (*J*_*i*_) in parallel by running the FIC algorithm. The FIC algorithm helps in establishing E-I balance in the whole brain. Under the same operant condition, we synthetically generate healthy FC from the model simulation. The obtained inhibitory synaptic weights (*J*_*i*_) are further considered as initial values for inhibitory plasticity in the following two conditions for the virtual lesion analysis.

In the middle row of Fig. 1b, the virtual lesion is introduced into the healthy SC, resulting in loss of E-I balance, i.e., a short-term loss of E-I balance due to lesion impact, as our second condition. Next, we simulate the virtual lesion model without FIC to capture altered FC for further comparison with healthy and other post-lesion conditions.

The lower row in Fig. 1b depicts the condition after the lesion when the FIC restores the E-I balance. At this condition, the adaptive nature of the brain re-adjusts the local inhibitory weights to restore the desired E-I balance. It allows the damaged brain to sustain at desired firing rate *∼* 4Hz. The virtual lesion model with the FIC is simulated to generate re-organized FC, which is compared against the altered FC obtained from the previous condition. While we numerically simulate the lesioned model, we store re-adjusted synaptic inhibitory weights (*J’*) and capture re-adjustment time (*RT*) during the reestablishment of local E-I balance in an individual area. Further, the curated *RT* and *J’* are correlated with the measured JC values corresponding to a lesion site. As we are more interested in finding changes in inhibitory synaptic weights, we calculate 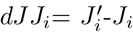 for all areas. Based on the two measured quantities (*RT* and *J’*), we estimate dynamically similar areas (DSAs) while observing the global homeostatic condition. We unveil the functional affinity for alternation and re-organization pattern of the brain after lesion by correlating SSAs, identified from anatomical measure *JC*, and DSAs from simulation parameters (*RT, dJJ*).

It is worth mentioning that the first condition is a one-time process, whereas the following two steps are repeated for different virtual lesion sites at the single subject level. All three steps have been repeated for individual subjects and further analyzed at the group level.

We have performed two levels of analysis, (i) anatomical level analysis and (ii) functional alteration/reorganization analysis. Significant changes between healthy and altered FCs and altered and re-organized FCs are captured by parametric test, an independent t-test analysis. Functional network properties (measuring network resilience, segregation, and integration) such as modularity, transitivity, global efficiency, and average characteristic path length are derived to evaluate functional alteration due to short-term loss of E-I balance (mimicking early lesion phase) and long-term functional re-organization in the post-lesion phase.

### Structural connectivity analysis

#### Identify SSAs using Jaccard coefficient

Weighted Jaccard coefficients (*JC*) are calculated from the individual subject’s healthy SC. High *JC* values imply a high structural similarity, whereas low values yield lesser similarity corresponding to an area of interest (could be a lesion center). A threshold is put on the obtained JC values corresponding to a lesion site to identify higher similar areas. The top 25% areas with higher *JC* values are considered as SSAs.

Figure 2a shows *JC* matrix obtained from a healthy SC (without lesion). Descending order distribution of *JC* values corresponding to the lesion at lPOPE is plotted in Fig. 2b. The top 25% similar areas are shown in blue bars and the rest in yellow. In Fig. 2c, only the top 25% similar areas (blue nodes) are plotted on the brain surface using BrianNet viewer. Node size implies *JC* values. The top 25% SSAs of lPOPE are lCMF, lRMF, lPTRI, lPREC, rSF, rCMF, lIP, lSP, rRMF, rPREC, lINS, lPCUN, lPCNT, rPTRI, rPOPE, written in descending order from the similarity indices. It is observed that the higher similar areas, e.g., rostral, caudate middle frontal cortex, parietal cortex, insular cortex, and primary and supplementary motor cortex, are found both in ipsilesional and contralesional hemispheres. SSAs corresponding to lPOPE for different subjects are shown in SI Appendix, Fig. S1a. The SSAs for other lesioned sites are shown in SI Appendix. Right POPE has SSAs such as lCMF, lPTRI, lPOPE, and lRMF in homotopic regions to the left hemisphere (SI Appendix, Fig. S1c), including rostral, caudal, precentral, and postcentral gyrus. SSAs of primary motor regions (left precentral gyrus, lPREC) are distributed in both hemispheres, including the caudal (l/rCMF), rostral (l/rRMF), frontal (l/rSF) cortex. Regions are also in the parietal (lIP, lSP, lPCUN) and insular (lINS) cortex for the left hemisphere (SI Appendix, Fig. S1e). The left lateral occipital (lLOCC), part of the visual cortex has SSAs mainly in the ipsilesional hemisphere ranging from the parietal lobe (lIP, lPCAL, lSP, lSMAR, lISTH) to the middle frontal lobe (lCMF) via temporal regions (lST, lMT, lTT, lFUS) and insular (lINS) cortex (SI Appendix, Fig. S1g). Other structural similar areas for different regions are tabulated in SI Appendix, Table S3.

**FIG. 2.**
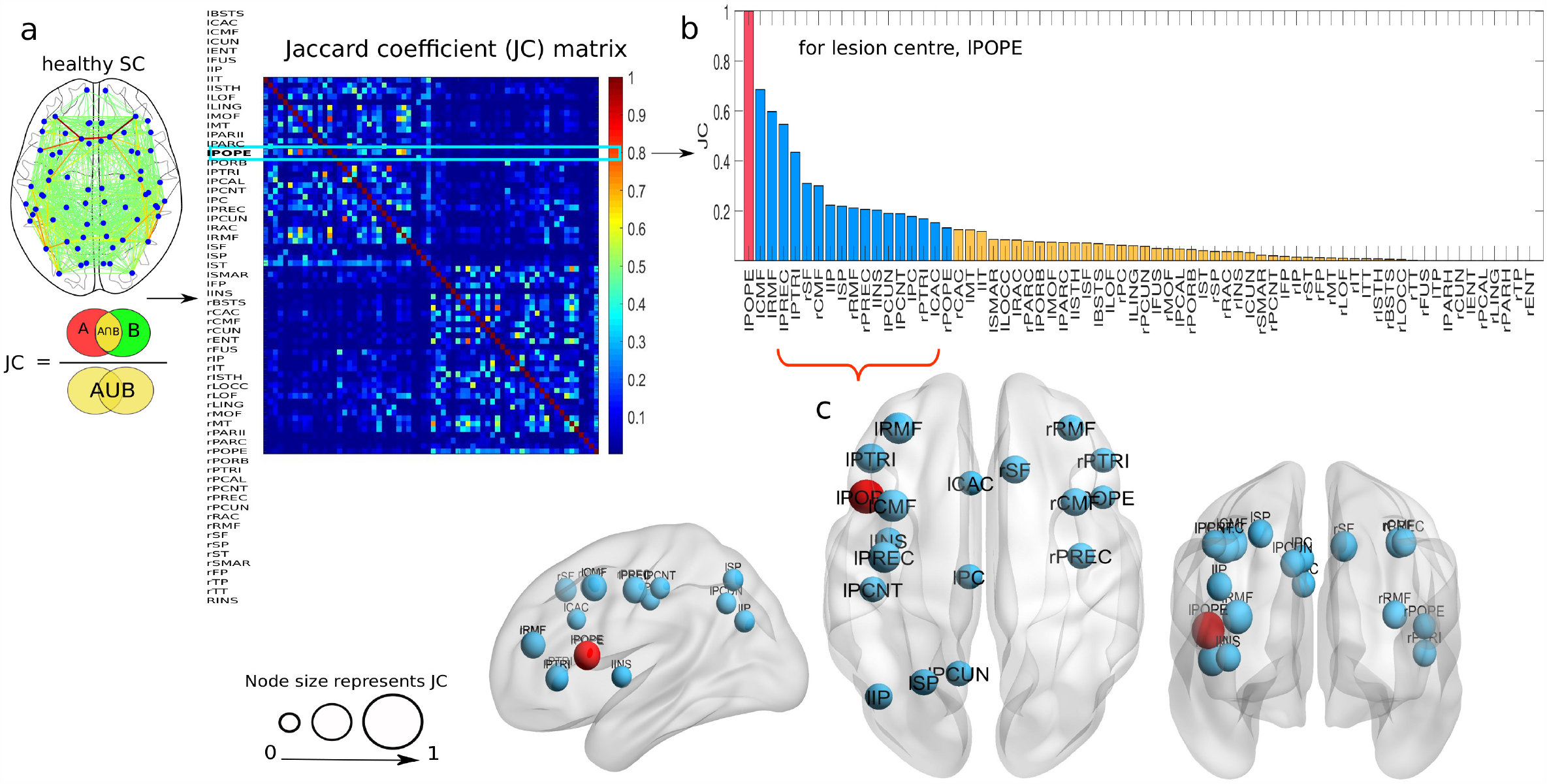
Similar structural areas (SSAs) identified by *JC* for a single subject. (a) *JC* is evaluated from healthy SC, where *A* and *B* are any two regions of interest. (b) Distribution of *JC* values, considering the lesion to be at left pars opercularies (lPOPE), are plotted in descending order. The red bar is the selected lesion site (lPOPE). The top 25% areas from the *JC* values are considered SSAs, shown in blue. The rest of the lower similar areas are shown in yellow bars. (c) Identified SSAs are shown in sagittal, axial, and coronal brain views. Node sizes represent *JC* values.

#### Correlation analysis between anatomical (JC) and dynamical (RT, dJJ) measures

The *JC* is derived from a healthy SC and used to test our hypothesis, whether the SSAs corresponding to a lesion site are essential in restoring E-I homeostasis in local regions and eventually within whole cortical systems. Model-based measurable parameters, re-adjustment time (*RT*), and change in inhibitory weights (*dJJ*) have been used to predict the dynamically similar areas (DSAs). We have simulated the virtual lesion model when the dynamical mean field (DMF) model is spatially connected via the virtually lesioned SC of a single subject. Area-wise distribution of *JC, RT*, and *dJJ* correspond to the lesion site at lPOPE is shown in Fig. 3a. A higher *RT* node value implies a more extended time required to reach the desired threshold in excitatory synaptic current for balancing the E-I ratio. A negative value of *dJJ* for an area implies a reduction in its inhibitory weight, i.e., a decrease in inhibition of that area. We take absolute *dJJ* for better visualization and description in our analysis. We find a positive association (*r* = 0.69) between *JC* and *RT*, which yields that SSAs take longer to readjust the E-I balance, as shown in Fig. 3b. Conversely, a positive correlation between *JC* and *dJJ*, fitted by a linear fitting model with *r* = 0.8, see Fig. 3c. It indicates that the SSAs have tuned their local inhibitory weights. We find a positive correlation between *RT* and *dJJ*, fitted by a linear regression model (*r* = 0.88); see Fig. 3d. Overall observations suggest that SSAs have a strong correlation with DSAs, which implies the SSAs take longer to modulate their inhibitory weights to settle the neural activity at the desired set point, i.e., balanced E-I ratio and target firing rate of *∼*4Hz.

**FIG. 3.**
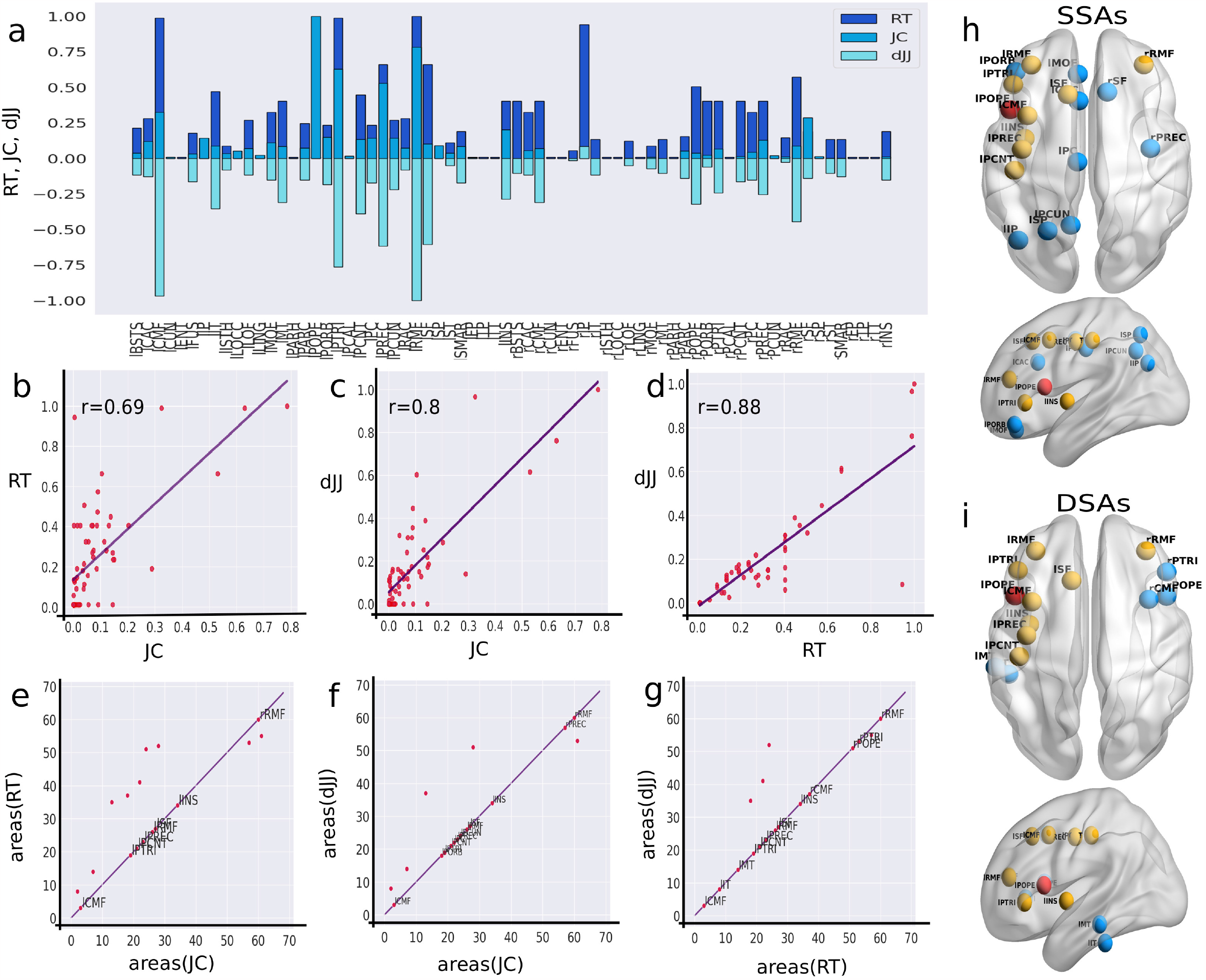
Correlation between anatomical and dynamical parameters corresponding to the lesion at lPOPE. (a) Area-wise distribution of *JC, RT*, and *dJJ* for a single subject. From the derived *JC* matrix on healthy SC, we only choose the *JC* values considering the lesion center to be at lPOPE. Next, we take absolute *dJJ* values for the rest of our analysis. (b-d) Scatter plots show the correlation between *JC*-*RT, JC*-*dJJ*, and *RT* -*dJJ*, respectively. Violet lines denote a positive correlation between any two given measures. Correlation values (*r*) are given at the top of each plot. (e-g) Areas of interest are identified considering a higher correlation between *JC*-*RT, JC*-*dJJ*, and *RT* -*dJJ*. The regions on the purple lines are common within the two measures. We choose DSAs from the overlapped areas in both *RT* and *dJJ*, i.e., the areas lie on the purple line in (g). (h) To compare between SSAs and DSAs, we repeat the identified SSAs here. (i) The estimated DSAs are shown in sagittal and coronal brain view. The red sphere is the lesion center, lPOPE. Common regions from SSAs and DSAs are shown in yellow and non-overlapping in blue.

Next, we aim to estimate the areas with a higher positive association measured from the correlation between *JC* and *RT*, or *JC* and *dJJ*. We have selected the regions that lie on the diagonal line (violet line) only, shown in Fig. 3e, which are common in both independent measures. The estimated regions, such as lCMF, lPTRI, lPCNT, lPREC, lRMF, lSF, lINS, and rRMF, are common in both SSAs and DSAs, with larger *JC* and *RT* values. Similarly, we identify regions from the correlations between *JC* and *dJJ*, such as lCMF, lPORB, lPTRI, lPCNT, lPREC, lPCUN, lRMF, lSF, lINS, rPREC, and rRMF, shown in Fig. 3f. Subsequently, we identify the predicted brain areas obtained from the substantial overlap between their *RT* and *dJJ* values. This approach identifies the following brain regions lCMF, lIT, lMT, lPTRI, lPCNT, lPREC, lRMF, lSF, lINS, rCMF, rPOPE, rPTRI, and rRMF; see Fig. 3g. The SSAs, selected from anatomical measure (*JC*), are shown in Fig. 3h. The DSAs, identified from dynamical measures (*RT* and *dJJ*), are plotted in Fig. 3i on the glass brain in sagittal and axial view. Yellow nodes in Figs. 3h,i are common in both SSAs and DSAs, where blue ones are the non-overlapping areas. The left pars opercularis (lPOPE) lesion site is shown in the red sphere. Interestingly, areas identified by the two independent analyses, i.e., SSAs and DSAs, have more than 60% overlapped regions corresponding to the lesion site, lPOPE. The estimated DSAs corresponding to lesion centers at lPOPE, rPOPE, lPREC, and lLOCC in different subjects are shown in SI Appendix, Figs. S1b, S1d, S1f, and S1h, respectively. Other DSAs for different lesion centers are tabulated in SI Appendix, Table S3. A strong correlation between *JC* and *RT* or *JC* and *dJJ* and high overlapping between SSAs and DSAs suggest that the predicted areas play a crucial role in re-establishing local E-I balance by calibrating their inhibitory weights and help sustain the target firing rate (4Hz) after the lesion occurrence.

Further, we sequentially introduce virtual lesions to all 68 areas. We investigate correlations between *JC* and dynamical measures (*RT, dJJ*) at the level of single subjects, as depicted in SI Appendix, Fig. S2. The correlation between *JC* and *RT* in Fig. S2a, and *JC* and *dJJ* in Fig. S2b is positive for different lesion centers. Except for a few regions, such as lENT, rENT, lPARH, rPARH, lFP, rFP, lTT, rTT, lTP, and rTP, other regions display largely weaker or negative correlations. The estimated SSAs and DSAs are displayed in SI Appendix, Figs. S3a,b. These ten nodes have less number of connections and nodal strength. Lower strength and degree of a node could be why their SSAs are not participating in E-I balance (SI Appendix, Figs. S3e,f).

#### Inter-subject and inter-hemispheric variability/similarity

Figure 4 shows the results for two subjects and lesion sites at two hemispheres. We describe the findings from the two independent analyses. The SSAs and DSAs corresponding to the lesion site lPOPE are repeated for one subject in Figs. 4a,b. Estimated common areas from both SSAs and DSAs are shown in yellow and non-overlapping in blue. The overlapping regions are lRMF, lPTRI, lCMF, lINS, lPREC, lPCNT, lSF, and rRMF, respectively.

**FIG. 4.**
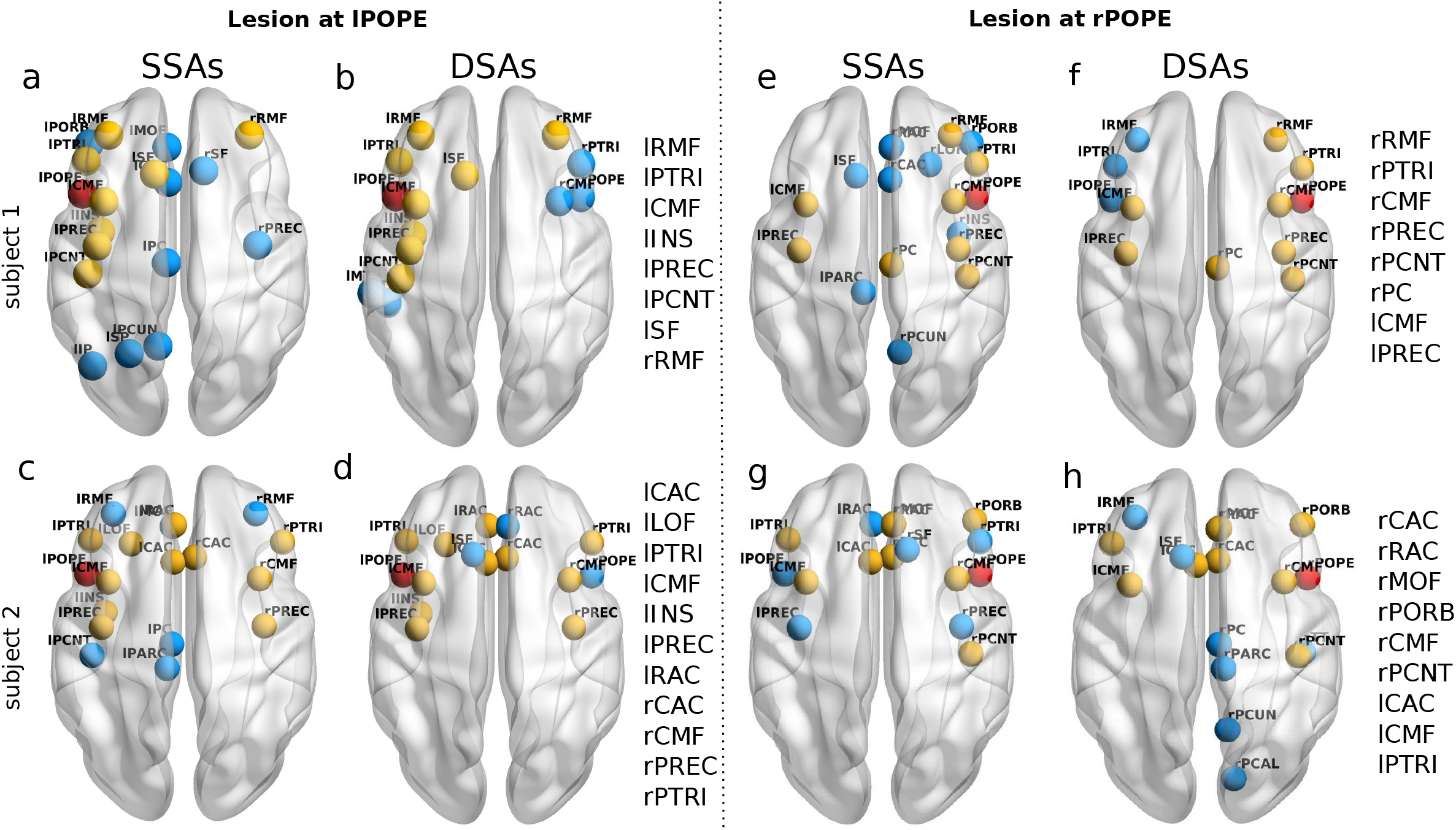
Inter-subject and inter-hemispheric variability/similarity in SSAs and DSAs. SSAs corresponding to lesion centers at left and right POPE (red spheres) are shown respectively in (a, c) and (e, g) for two subjects. DSAs are shown respectively in (b,d) and (f,h) for those two subjects. Common regions from SSAs and DSAs are shown in yellow spheres. Abbreviations are written on each subplot’s left side. Overlapping areas in SSAs and DSAs are yellow, and non-overlapping are blue.

Inter-subject variability is depicted for another subject in Figs. 4c and 4d. The overlapping regions for this subject are lCAC, lLOF, lPTRI, lCMF, lINS, lPREC, lRAC, rCAC, rCMF, rPREC, rPTRI, displayed in yellow in Figs 4c,d. Although the lesion centers are similar for both subjects; still, the identified SSAs and DSAs are different in the two subjects; compare Figs. 4a and 4c, or Figs. 4b and 4d. Variability in SSAs arises from individual subjects’ structural/anatomical differences in brain connectivity and manifest individual-specific SC-FC correlations.

However, it is interesting to note that the areas predicted for lesion recovery by *JC* are similar to the regions predicted by the dynamical measures (*RT* -*dJJ*) in individual subjects (see Figs. 4a,b or, Figs. 4c,d), despite inter-subject structural differences. While comparing the two Figs. 4b and 4d, the estimated DSAs lie in the anterior cingulate cortex (l/rRAC, l/rCAC) for subject-2 (Fig. 4d), whereas no nodes from the anterior cingulate cortex are found for subject-1 (Fig. 4b). Patterns of subject-dependent variability/similarity in the estimated DSAs and SSAs are consistent for different lesion locations and tested for several subjects (SI Appendix, Fig. S1).

Further, we tested our hypothesis for another lesion site in the right hemisphere, say right pars opercularis (rPOPE), for the two subjects. We find consistent variability patterns in the results, see Figs. 4e,g or, Figs. 4f,h, and similarity in identified regions, comparison between the SSAs and DSAs in Figs. 4e,f or Figs. 4g,h. The overlapped regions from the SSAs and DSAs are shown in yellow, and the non-overlapping in blue.

#### Group-level analysis on SSAs and DSAs

A group-level analysis is performed over all 49 subjects to find the probability of the appearance of a predicted SSA or DSA. We determine the probability of appearance (PA) of an area as a ratio between the number of an SSA (or DSA) that appeared within all the subjects and the total number of subjects as,

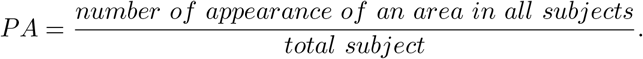

The value *PA*=1 corresponding to an SSA (or DSA) implies that it appeared in all the subjects. The values and distribution of the PA for SSAs corresponding to lPOPE are shown in Figs. 5a, and 5b, respectively, and the PA values for DSAs are plotted in Figs. 5c,d. Yellow nodes and bars stand for higher PA values, and pink shade implies lesser PA values, indicated by the color bar. It is observed that the SSAs corresponding to the lesion center at lPOPE, such as lCMF, lPTRI, lPREC, lRMF, and lINS, are found in all 49 subjects, see Figs. 5a,b. Other SSAs, including the postcentral gyrus, precuneus, posterior cingulate, and contra lesional frontal regions, are found in more than 90% of the subjects. SSAs in the right hemisphere, e.g., rPOPE and rPTRI, are found in more than 50% of subjects. Similarly, the DSAs, including lCMF, lPTRI, lPCNT, and lRMF, are found in almost all the subjects. We test the consistency and robustness of our results for other lesion locations; please check SI Appendix, Figs. S3a,b, and Figs. S3e,f.

**FIG. 5.**
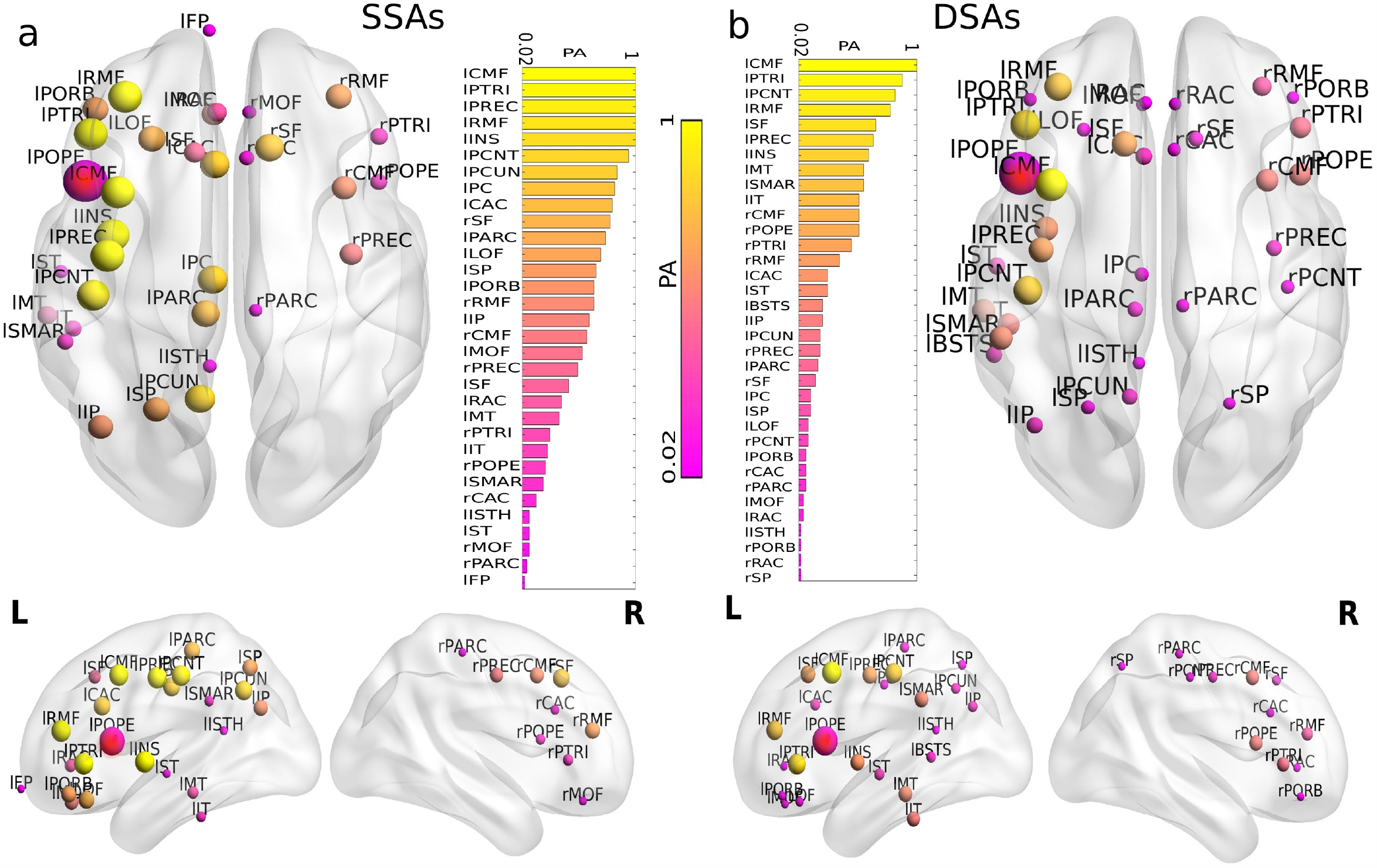
Group-level analysis on SSAs and DSAs corresponding to lPOPE. Probability of appearance (PA) of nodes identified by structural and dynamical measures over subjects corresponding to the lesion at lPOPE are illustrated. (a) Axial, sagittal (left and right) view of the brain for SSAs over all subjects and the distribution of PA in descending order for SSAs. (b) Descending order distribution of PA for DSAs, axial and sagittal (left and right) view of the brain for DSAs over all subjects. Yellow areas have higher PA yielding, appearing as SSA and DSA over all subjects. The pink areas appeared less in both SSAs and DSAs. The red node is the lesion site, lPOPE.

Although the identification of SSA based on anatomical property is entirely independent of the estimation of DSA, both these proposed methods can crucially identify and predict similar compensatory candidate brain regions likely initiating the post-lesion recovery process. The higher similarity between SSAs and DSAs signifies that the similar structured areas corresponding to a possible lesion site have modulated their local inhibitory weights and participated in restoring local and global homeostatic E/I balance.

### Functional alteration and re-organization after lesion

We have used statistical tools to investigate how anatomical perturbation to a node affects the global functional organization. The impact of lesion on FC has been categorized into two parts: i) alteration and ii) reorganization. Simulated FC is obtained from the spatiotemporal BOLD signals using pairwise Pearson correlation. Considering three conditions for each subject, we have synthetically generated healthy FCs, altered FCs when E-I balance is lost and re-organized FCs when the E-I balance is restored. Statistical comparison between any two conditions, e.g., healthy vs. altered FCs, and altered vs. re-organized FCs, respectively, determines significant differences between generated FCs. First, we calculate the z-score of individual FCs for the three conditions. Next, we perform paired sample t-tests to designate the significant global changes in the healthy normative pattern, i.e., deviation from the healthy FC into the altered FC that depicts the direct impact of lesion on collective dynamics. Similarly, after the global restoration of E-I homeostatic balance, the post-lesion functional reconfiguration pattern is investigated by comparing the altered and re-organized FC. The ROI-wise paired t-test. is performed for each element in the FC matrix between two conditions, e.g., healthy and altered FC group and altered and re-organized FCs, for all subjects. False discovery rate (FDR) is corrected over the obtained *p*-values from the t-test.

Figure 6 shows ROI-wise FC analysis between two conditions, e.g., healthy-altered FC and altered-reorganized FC, considering lesion center lPOPE. Subject-wise model-generated healthy FC, altered FC, and reorganized FC are shown in Figs. 6(a-c), respectively. ROI-wise *t*-statistic to find significant changes in the weights between healthy and altered, as well as altered and re-organized FCs, are shown in Figs. 6d and 6e, respectively. Upper triangular elements in Figs. 6d,e represent the *t*-statistics corresponding to changed weights between any given pair of regions. Lower triangular values in Figs. 6d,e is obtained by putting a threshold on *p*-values (*p <* 0.005).

**FIG. 6.**
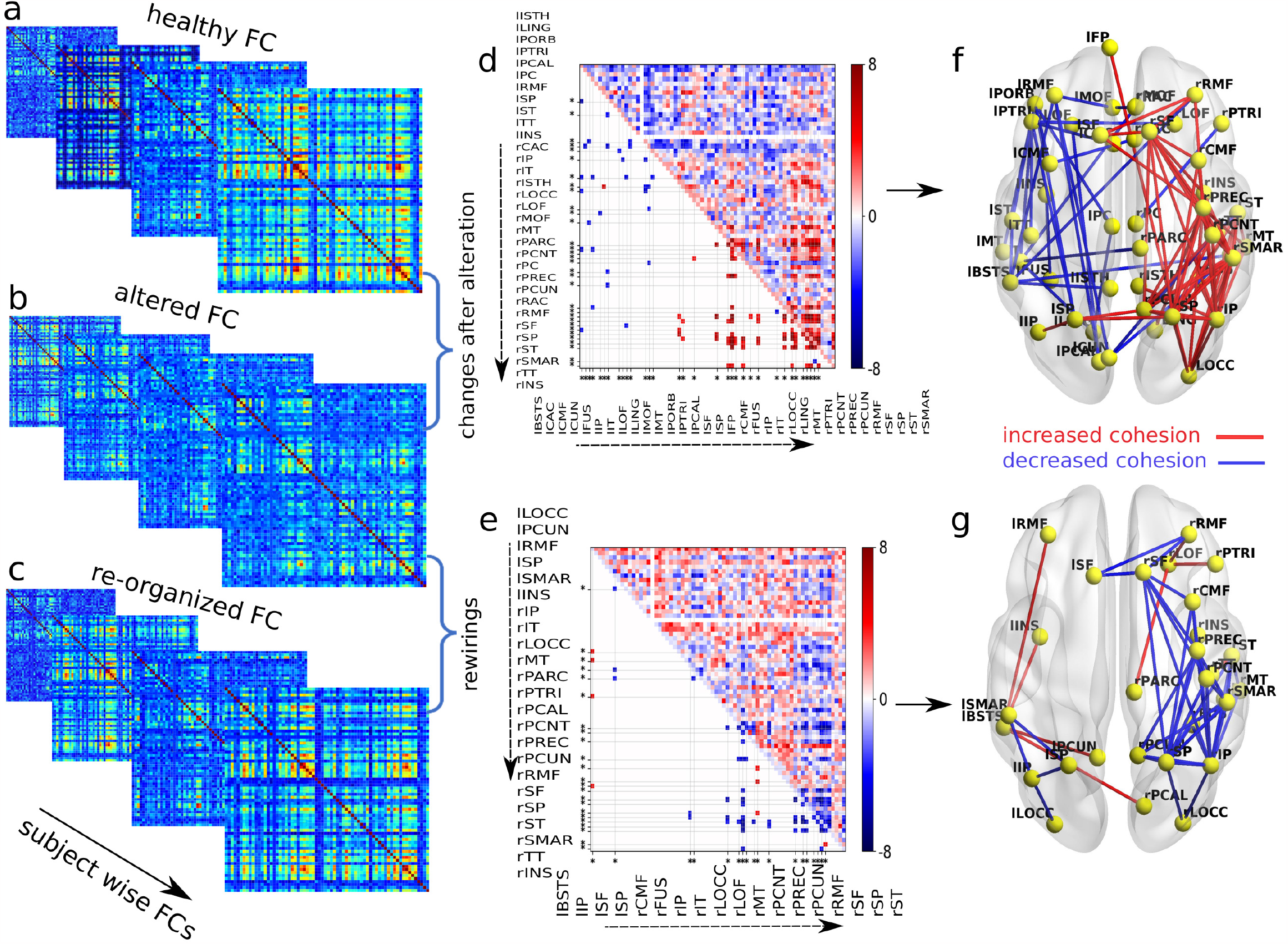
ROI-wise FC analysis. (a-c) Model-generated healthy, altered, and re-organized FCs are placed in three rows. The virtual lesion is introduced at lPOPE. ROI-wise changes and rewiring between two conditions over the subjects are measured using paired t-tests. ROI-wise *t*-statistics are displayed in the upper triangular matrices, comparing healthy and altered FCs in (d) and altered and re-organized FCs in (e). Lower triangular entries are the significant *t*-statistic, putting a threshold on the *p*-values (*p <* 0.05, FDR corrected). The color bar indicates the *t*-stat value. Stars at the bottom and left sides of the matrices are significantly changed regions. Glass brain plots in (f) and (g) show, respectively, the significantly changed and rewired links with related regions. Red and blue edges represent the significant increase and decrease in *t*-stat values. (f) Due to lost homeostatic balance, cortical cohesion is reduced in ipsilesional and increased in contralesional. In contrast, (g) spatiotemporal correlation is enhanced in the ipsilesional and attenuated in the contralesional hemisphere after restoring the E-I balance.

It can be noticed that cortical cohesion is significantly decreased in the ipsilesional hemisphere but increased in the contralesional hemisphere, shown by the red and blue lines in Fig. 6f. Furthermore, when the E-I balance is globally restored, we observe a significant increase in synchrony in the ipsilesional hemisphere and a decrease in the contralesional hemisphere, shown in red and blue in Fig. 6g. Results for other lesion centers are shown in SI Appendix, Fig. S3.

## III. DISCUSSION

In this work, we have proposed two independent measures SSAs (anatomically self-similar areas) and DSAs (dynamically self-similar areas), using virtual lesion modeling in predicting candidate brain regions that initiate the post-lesion recovery by reestablishing local and global E/I balance. We have demonstrated the compensatory role of SSAs corresponding to a lesion site/center in restoring E-I balance in the local areas and across the whole brain mediated by the negative feedback form of plasticity. Resetting homeostatic balance results in postlesion functional re-organization within the surviving cortical areas. The homeostasis mechanisms, governed by the Feedback Inhibition Control(FIC) [41], are deployed to restore E-I balance within the virtually lesioned brain regions at the group and subject-specific level. We measure the simulation time (called re-adjustment time, *RT*) taken by each region to re-adjust their local inhibitory weights (*J’*) during the re-establishment of global homeostatic balance after lesion. Our hypothesis of predicting compensatory brain regions is based on structural and functional equivalence. Further, the observed compensatory utilization of the hemisphere is supported strongly based on SC analysis and FC alteration/rewiring.

### Observations from SC analyses

Findings from the structural analysis are divided into two parts, (i) identification of SSAs and (ii) estimation of DSAs. From the anatomically constrained dynamical model simulation, we have estimated the DSAs, further correlated against the SSAs. Obtaining the correlation between SSAs and DSAs has helped to test our proposed hypothesis. Despite the diversity in inter-individual structural topology, the two independent methods (SSAs and DSAs) provide computational machinery for predicting common brain areas with 60%-70% overlaps. To check the probability of an estimated region being a potential candidate area for functional recovery, we have measured the probability of appearance (PA) within all 49 subjects in our data. A Higher PA corresponding to a predicted area indicates a high chance of being a candidate in all subjects and a high probability of participating in compensation for the damaged brain.

From the derived PA, the SSAs corresponding to the lesion at left POPE (associated with language processing) is identified as caudal middle frontal gyrus (CMF), rostral middle frontal gyrus (RMF), inferior frontal gyrus (IFG), primary and secondary motor areas (PREC, PCNT), and insular cortex (INS) in the ipsilesional hemisphere. Contralesional homologous regions are rPOPE and rPTRI. Dynamically similar areas for left POPE are found in both ispi- and contra-lesional hemispheres, including CMF, RMF, PTRI, PREC, and PCNT. Predicted areas belonging to the SSAs and DSAs in both ipsi- and contra-lesional regions suggest a CRUH. The predicted areas are independent of the subject’s age and gender. The identified SSAs from both hemispheres are also reported in earlier studies as essential candidates for complete language recovery after lesion. For example, the recruitment of perilesional tissue [46], as well as contralesional areas [32, 47–51], and participation of the homologous regions [52–54] in association with language recovery are well documented largely concurs with our findings. Besides, increased activation of right lesion-homolog inferior frontal gyrus (IFG) has been reported in a subset of patient groups [54].

For a lesion site at the primary motor region, left postcentral gyrus (lPREC), we have predicted candidate areas as CMF, RMF, pars triangularis, superior parietal, superior frontal gyrus, and precuneus in the ipsilesional hemisphere. Moreover, Superior frontal and rostral middle frontal regions from the contralateral hemisphere are found in almost all subjects. Our observations align well with previous findings on ‘motor lesion re-organization in ipsilateral premotor cortex [55, 56], recruitment of contralesional motor areas [57–59]. The areas reported by neurological studies found to be crucial in motor function recovery [30]. Other motor-related brain regions such as supplementary motor area (SMA), dorsolateral premotor cortex (PMC) and cingulate motor areas (CMA), and insular cortex [31] provides necessary compensation to improve motor performance [60] as documented by previous findings. Also, found to be critical for functional reorganization in the motor recovery [61, 62] and matches completely with the predicted brain regions based on our proposed computational framework.

In contrast, few regions such as entorhinal (ENT), parahippocampal (PARH), frontal pole (FP), temporal pole (TP), and traverse temporal (TT) regions have displayed significantly less correlation between *JC* and *RT* (SSAs and DSAs). The weaker association, in this case, may arise due to their sparse connectivity and less anatomical strength. Lesions in these areas have a much lesser impact on overall homeostasis. Thus, the damage to these brain regions may restore the lost E-I balance with comparatively minimal effort. In the above scenario, the adjacent brain regions participated in resetting local and global homeostasis other than SSAs (or DSAs), suggesting local wiring specificity and proximity could be key to initiating the neural compensatory process.

To this end, we mention two major aspects of our findings. First, a large overlapping brain region predicted by SSAs and DSAs signifies structurally similar regions primarily participate in the dynamical re-organization process. These areas reset local E-I balance after lesion by modulating their inhibitory weights, thus, displaying the constructive role of SSAs on functional network recovery. It can be concluded that a higher correlation between these two independent methods (SSAs and DSAs) arises from the interplay between the structural property and the local inhibitory weights responsible for emergent globally coordinated dynamics. Specifically, an emerging local re-adjustment of inhibitory weights mediates self-organized global brain dynamics during homeostasis. The second aspect of the findings is that the SSAs are identified from the healthy subject’s SC analysis. In contrast, the DSAs are estimated after introducing virtual lesions in the SC matrix. Thus, a methodological advantage is that even if, in the clinical phase, healthy FC is unavailable for a particular patient, the DSAs can be employed as a tool in patient-specific FC to identify candidate compensatory brain areas.

### The compensatory mechanism

In addition, simulation of the virtual lesion model has helped to acquire insight into the dynamic origin of the post-lesion compensatory mechanism. We find that the regions from both ipsi- and contra-lesion, with higher structural similarity to the lesion site, took an extended time to modulate their local inhibitory weights. Besides, the SSAs, from both hemispheres have reduced regional inhibition to balance decreased excitation due to degradation in the excitatory synaptic drive, thus balancing the overall E-I ratio and sustaining the target firing rate of 4Hz. As documented earlier, homeostatic plasticity, a mechanism of up-and-down regulation of both the presynaptic release of and the postsynaptic response to neurotransmitters, is essential to maintain a stable set point and near-normal brain condition [26]. Here, the primarily recruited SSAs from the two hemispheres suggest the dominant role of structural similarity and rewiring proximity of compensation-related utilization of hemispheres, which have guided homeostatic plasticity-driven compensatory mechanisms in re-organizing post-lesion functional brain network recovery.

### Observations from FC analyses

Previous studies have reported that changes in excitability affect the local excitatory-inhibitory (E-I) balance of the lesion site and distant cortical networks [63], known as diaschisis [64], suggesting remote disruptions in FC following lesion impact. These studies have hypothesized mechanisms underlying neuronal remodeling in the perilesional area and contralesional hemisphere after motor cortex infarcts and summarized evidence from previous studies based on analysis of electrophysiological data that demonstrated brain-wide alterations in functional connectivity in both hemispheres, well beyond the infarcted area.[65, 66]. Our findings based on FC analyses depict reshaping in the coordinated cortical cohesion, which is not limited to the ipsilesional hemisphere but also progresses distant from the damaged area into the contra-lesional hemisphere showing nonlocal effects and completely aligned with the experimental findings from human and animal studies depicting brain regions implicated during the post-lesion functional recovery process. In our study, we have also observed that an early impact of structural damage results in a specific signature, the reduction in spatiotemporal cortical coherence in the ipsi-lesional hemisphere. On the contrary, the spatiotemporal coherence increases in the contra-lesional hemisphere. In the emergent FC, at E-I balanced state after lesion occurrence, we observed increased synchronous neural activity in the ipsi-lesional site, while a decreased synchronous neural activity in the contra-lesional hemispheres. This seesaw effect in the opposing hemisphere to the lesion center aided functional restoration. The re-organization pattern in the ipsi- and contra-lesional hemispheres is similar in all subjects and for different lesion sites heralding the robustness and consistency of the findings reported here. In response to structural damage, we find that the cortical plasticity mechanism related to E-I homeostasis facilitates FC rewiring in the contra-lesional hemispheres, similar to the previous key observations in [24]. Our proposed computational mechanisms following lesion could be similar to re-organization in pre-infracted to and distant regions from the lesion site that may trigger large-scale remodeling of the cortical networks to compensate for post-lesion deficits [6].

In addition, we have reconfirmed our observations using graph-theoretical properties of FC [1, 67, 68] such as transitivity, path length, modularity, and global efficiency, which are interpreted in terms of lesion impacts, and FC recovery. Due to lesions, the biological perturbation reshapes the healthy normative pattern in FC. The loss of inter-area excitatory synapses predominantly affects coordinated neural dynamics, which can be observed from the segregated functional network captured by the increased modularity and decreased global efficiency. However, the damaged brain tries to adapt to the global changes in functions caused by the homeostatic imbalance across the whole brain. Our *in − silico* investigation suggests that the adaptive mechanism compensates for the lost E-I homeostasis, primarily driven by the modulation of inhibition at the local level, mainly within the SSAs (or DSAs) identified in this work.

Gradual resetting of neural activity to regain the near-normal function is confirmed by the decreased modularity and increased global efficiency concerning the altered FCs. The FC integration has compensated segregation of FC after the damage to underlying structural connections. This further resets healthy dynamical repertoire driven by the negative feedback-mediated form of plasticity. Further, it is well explained in earlier studies [10, 69] that the global homeostasis is balanced by increasing excitability in the areas near and distant to the lesion center, suggesting a direct correlation between the E-I balance and global cortical dynamics, which can be one crucial aspect in proper lesion recovery.

### Summary of the findings

Previous studies [2, 70] have explored the critical role of network topology in the context of the lesion. However, whether compensatory brain regions could be predicted based on identifying topologically self-similar and dynamical self-similar areas (an equivalence principle) is largely unknown. Here, we propose this equivalences principle by introducing two new tools grounded on topological and dynamical self-similarity, which allow us to predict compensatory brain areas and mechanisms during post-lesion recovery. Our proposed theoretical framework identifies overlapping compensatory brain regions during the early and post-lesion recovery phase across all subjects using two independent methods and subjectwise variability. Our results provide the first unified framework behind observing a variety of compensatory brain regions identified by earlier lesion recovery studies. The cause and consequence of those identified areas remain a large knowledge gap in the neuroscience literature. We also report that SSAs are more robust and reliable than other network properties widely used in the literature (e.g., clustering coefficient, participation coefficient, node weight) when harnessing region-specific roles in reshaping near-normal functional brain connectivity and dynamics to identify the post-lesion recovery process. The DSAs provide a broader scope for investigating the post-lesion period when SSA is undetermined. The predicted areas (SSAa or DSAs) are independent of the subject’s age and gender. The key compensatory mechanism demonstrated here suggests a CRUH in the emergence of post-lesion coordinated cortical cohesion. Most importantly, the proposed theoretical methods are general and can be applied to broader lesion categories.

### Limitations

There are also limitations of this study. (i) A precise mapping between the accurate lesion biological time scale and simulation time is still being determined. Therefore, we cannot predict the time scale of the actual recovery process. Mapping real-time-scale with simulation time can bring us closer to uncovering the true recovery mechanism and will have excellent translational value. We are currently investigating this mapping in another research work and out of the scope of this study. (ii) The Feedback Inhibitory Control (FIC) mediates the homeostasis mechanism. However, other feedback-mediated mechanisms may be incorporated into the model to verify increased excitability due to lesions in individual subjects, which this work does not sufficiently explore. (iii) We did not incorporate the effect of lesion volume in this study. The amount of lesion volume and spread are essential factors in FC re-organization and recovery, which are not addressed here. (iv) Directed FC also holds the key to understanding how information flow alters following lesion and during recovery, which future studies may explore, (v) Finally, how FC is reshaped based on longitudinal data can validate present findings in a more nuanced fashion and establish a stronger link between lesion recovery and functional re-organization elucidating region-specific roles.

### Conclusion and future aspects

In conclusion, we envision a novel method to identify potential candidate areas responsible for resetting E-I homeostasis as possible compensatory mechanisms resulting in near-normal functional brain network recovery. A fundamental open question in the literature is how the non-lesioned brain adapts to the post-injury functional recovery process and whether those areas could be predicted using a systematic theoretical framework. Although the lesion recovery process may be complex, the current study provides a general framework elucidating that brain recovery involves the utilization of an equivalence principle based on structural and dynamic similarity to tackle a wide variety of lesions. Future studies could use controllability theory to narrow the DSA (SSA) estimation into a specific region on the directed FC network. Those studies could further pinpoint whether a DSA (SSA) driven information flow pattern exists in the surviving cortex. Furthermore, how do the candidate areas help information fidelity during the lesion recovery?

## IV. MATERIALS AND METHODS

### Participants

Resting state MRI data from 49 healthy subjects (31 females), ages ranging from 18 to 80 years (mean age 41.55 *±* 18.44 years), have been collected at Berlin Center for Advanced Imaging, Charit’
se University Medicine, Berlin, Germany [44].

### Anatomical connectivity

Resting state MRI, diffusion-weighted MRI, and functional MRI are performed using a 3 Tesla Siemens Tim Trio MR scanner and a 12-channel Siemens head coil. Detailed information on data acquisition parameters is found in [44]. We did not process the raw data. The data was pre-processed, and structural connectome was generated previously, using the pipeline by Schirner et al. [44]. Cortical grey matter parcellation of 34 regions of interest(ROI) in each hemisphere is considered following Desikan-Killiany parcellation [45]. The SI Appendix, Table S1 shows all the regions of interest (ROIs) with abbreviations.

### Empirical functional connectivity

Participants are subjected to a functional MRI scan in eyes-closed awake resting-state condition. The restingstate BOLD activity is recorded for 22 minutes (TR=2 sec). Pre-processing steps are given in SI Appendix. After pre-processing, aggregated BOLD time series of each region is z-transformed. The pairwise Pearson correlation coefficient is computed to obtain each subject’s restingstate functional connectivity (rsFC) matrix.

### Definitions and descriptions

Defining and describing the terminologies related to the study is worthwhile before drawing the pipeline and workflow. The definitions of the Jaccard coefficient, virtual lesion, DMF model, virtual lesion model, re-adjusted inhibitory weights, and time to reach E-I balance are presented below:

### Jaccard coefficient (*JC*)

Jaccard coefficient (*JC*) measures the pairwise correlation between any two brain areas, defined by the ratio between the sum of their common neighbors’ weights and the total weights of their neighbors [71]. The weighted Jaccard coefficient is expressed as 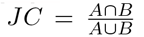, where *A* and *B* are the neighbors (specifically, the edge strengths with neighbors) of any two brain areas from the anatomical connectome of healthy subjects. The JC is measured with an unperturbed structural connectivity (SC) matrix before introducing a virtual lesion in the SC matrix at the level of an individual subject. The nodes with higher JC values corresponding to a lesion site share a similar topological property with the lesioned location mentioned as SSAs.

### Dynamic mean field (DMF) model

We use a reduced dynamic mean field (DMF) model [72] to engender lesion effects. The DMF approximates a spiking network model [41, 73] consisting of populations of excitatory and inhibitory neurons with excitatory NMDA synapses and inhibitory GABA synapses. DMF is described by a set of coupled nonlinear stochastic differential equations given below,

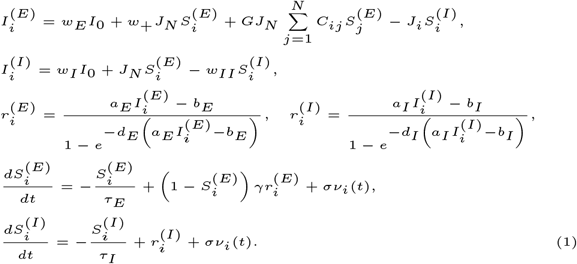

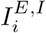 is the input current to area *i* and superscripts represent excitatory (*E*) and inhibitory(*I*) populations in that area. 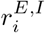 is the population firing rate of excitatory or inhibitory populations of area *i*. 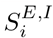 is the average excitatory (inhibitory) synaptic gating variable of area *i. I*_0_ is the effective external input scaled by *w*_*E*_ and *w*_*I*_ for excitatory and inhibitory populations. *w*_+_ is the local excitatory recurrence, *J*_*N*_ is the excitatory synaptic coupling, and *J*_*i*_ is the local feedback inhibitory synaptic coupling. *w*_*II*_ is the local inhibitory recurrence. *C*_*ij*_ is the *i, j*^*th*^ entry in the SC matrix, obtained from diffusion imaging, that scales the long-range excitatory currents between *j*^*th*^ and *i*^*th*^ regions. *G* represents global coupling strength which scales long-range excitatory connections. To find optimal *G*, the DMF model is simulated for different values of *G*. The optimal value of *G* is chosen based on the highest correlation between empirical and simulated FC when the excitatory firing rate sustains at *∼*4Hz within all brain regions [43, 74]. Euler’s method, with a step size of 1 ms, has been used to generate the synaptic activity of each area. The model is simulated for 10 minutes, where the first 2 minutes of transients are discarded. Parameter values are given in SI Appendix, Table S2.

### Feedback Inhibition Control

FIC algorithm, proposed by Deco et al. [41] is a recursive process to establish and maintain E-I balance in individuals and across all cortical subunits. The detailed FIC steps are found in the supplementary material. We simulated the model with the FIC algorithm for 10 *sec* time windows.

### Virtual lesion

The virtual focal lesion is introduced into an individual subject’s SC by targeted node removal. All connections to and from the focal lesioned site have been set to zero in the SC matrix.

### Virtual lesion model

We put the DMF model on top of a virtually lesioned SC of a single subject, labeled as a virtual lesion model. Individual node dynamics are governed by the stochastic DMF model spatially coupled via lesioned SC matrix. In principle, any other form of lesioned SC (real, virtual), if incorporated into the dynamical model, can produce similar effects of lesion depending on the lesion type, location, and extent.

### Modulated local inhibitory weight (*J’*)

The modulated local inhibitory synaptic weights 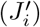 at individual nodes, in case of a lesion are compared against the weights estimated in a healthy brain (*J*_*i*_). The change in inhibitory synaptic weight is calculated as 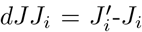 for all areas, *i*.

### Re-adjustment time (*RT*)

In the process of FIC, brain areas take different simulation times to reach the desired firing rate and balanced E-I homeostasis. The separate area’s re-adjustment time (*RT*) is estimated by tracking those simulated time windows by re-setting the E-I balances in all regions until the entire brain reaches the targeted balanced state. Each region modulates the local inhibitory-to-excitatory synaptic weights while restoring the E-I balance across the whole brain. During the FIC process, we track the elapsed time *RT*_*i*_ and *J’* for all regions, measuring units *sec* and *mMol*, respectively. Further, we use the obtained *RT* and *dJJ* to identify DSAs..

### Supporting Information (SI)

SI Appendix contains Tables S1-S3, Figs. S1-S7, details of the empirical data, DMF model description, and SI references. Table S1 shows model parameter descriptions and default values. Table S2 contains region names and abbreviations. Table S3 contains the list of SSAs and DSAs for different lesion centers. Predicted SSAs and DSAs for ten subjects in Fig. S1. Association be-tween structural and dynamical measures for different lesion centers over all subjects in Fig. S2. Details of the regions with lower connectivity are in Fig. S3. Figure S4 shows results for group-level analysis on SSAs and DSAs. ROI-wise FC analysis over all subjects for different lesion centers is shown in Fig. S5. Correlation between JC and other network properties of SC in Fig. S6. Figure S7 depicts FC properties under three conditions.

## ACKNOWLEDGMENTS

SS is supported by NPDF, SERB-DST, India, Award ID: PDF/2021/000585. DR and AB acknowledge NBRC Flagship program, DBT, India, Award ID: BT/MED-III/NBRC/Flagship/Flagship2019. AB is supported by the Ministry of Youth Affairs and Sports, India, Award ID:F.NO.K-15015/42/2018/SP-V. DR is supported by Ramalingaswami Fellowship, DBT, India, Award ID: BT/RLF/Re-entry/07/2014

## Supplementary Information for

### Empirical data

#### Participants

MRI and resting-state functional MRI data from 49 healthy subjects (31 females), ages ranging from 18 to 80 years (mean age: 41.55 years; standard deviation: 18.44 years), have been collected at Berlin Center for Advanced Imaging, Charité University Medicine, Berlin, Germany (1). The participants are healthy, and no history of neurologic or psychiatric conditions was reported in (1). All participants gave written informed consent to the group (1), and the study was performed under the compliance of laws and guidelines approved by the ethics committee of Charité University, Berlin, Germany.

#### Anatomical connectivity

Resting state MRI, diffusion-weighted MRI, and functional MRI are performed using a 3 Tesla Siemens Tim Trio MR scanner and a 12-channel Siemens head coil. Detailed information on data acquisition parameters is found in (1). We did not process the raw data. The data was pre-processed, and structural connectome was generated previously, using the pipeline by Schirner et al. (1). Cortical grey matter parcellation of 34 regions of interest(ROI) in each hemisphere is considered following Desikan-Killiany parcellation (2). The SI Appendix, Table S1, shows all the regions of interest (ROIs) with abbreviations.

#### Empirical functional connectivity

Participants are subjected to a functional MRI scan in eyes-closed awake restingstate condition. The resting-state BOLD activity is recorded for 22 minutes (TR=2 sec). Pre-processing steps are given in SI Appendix. After pre-processing, aggregated BOLD time series of each region is z-transformed. The pairwise Pearson correlation coefficient is computed for each subject’s resting-state functional connectivity (rs-FC) matrix.

#### Preprocessing of empirical data

##### Structural connectivity

Each subject’s empirical structural connectivity (SC) was generated using the pipeline described by Schiner et al. (1). Main pre-processing steps for T1 anatomical images involved skull stripping, removal of non-brain tissue, brain mask generation, cortical reconstruction, motion correction, intensity normalization, WM, subcortical segmentation, cortical tessellation generating GM-WM and GM-pia interface surface-triangulations and probabilistic atlas based cortical and subcortical parcellation. Cortical grey matter parcellation of 34 regions of interest(ROI) in each hemisphere was undertaken following Desikan-Killiany parcellation (2). The probabilistic tractography algorithm estimated the connection strength (a value ranging from 0 to 1) between each pair of ROIs. SC matrices were generated from each subjecTs MRI data and then summed element-wise to obtain an averaged SC matrix. The connection of a region to itself was set to 0 in the SC matrix for the simulations. motion correction and eddy current correction (ECC), the b0 image is linearly registered to the subject’s anatomical T1-weighted image, and the resulting registration rule is used to transform the high-resolution mask volumes from the anatomical space to the subject’s diffusion space. MRTrix has been used to extract gradient vectors and values (b-table). DW-MRI data were pre-processed using FREESURFER. The pre-processing steps for the diffusion MRI data were eddy current and motion correction with re-orientation of b-vectors (b-zero image was linearly registered to the subject’s anatomical T1-weighted image). Then, fiber-response function estimation has been done. The fiber orientation distribution function (fODF) for each image voxel has been computed based on constrained spherical deconvolution (CSD) in MRTrix. Structural connectome is the count of tracks between any given pair of ROIs. SC is normalized and symmetric.

##### Functional connectivity

Each subject’s empirical functional connectivity (FC) was computed using the pipeline described by Schiner et al. (1). To generate the functional connectivity (FC) matrices, pre-processing steps are as follows: 1) raw fMRI DICOM files were converted into a single 4D Nifti image file. 2) FSL’s FEAT pipeline is used to perform the following operations: a) deleting the first five images of the series to exclude possible saturation effects in the images, b) high-pass temporal filtering (100 seconds high-pass filter), c) motion correction, d) brain extraction and e) a 6 DOF linear registration to the MNI space. 3) BOLD signals are registered to the subject’s T1-weighted images and parcellated according to FREESURFER’s cortical segmentation (Desikan-Killiany (DK) atlas (2)). 4) The inverted mapping rule mapped Anatomical segmentation onto the functional space. 5) Average BOLD signal time series for each ROI were generated by computing the mean of all voxel time series of each region. 6) From the region-wise aggregated BOLD data, FC matrices were computed within MATLAB using pairwise mutual information (on z-transformed data), and Pearson’s linear correlation coefficient as FC metrics.

#### Dynamic mean field (DMF) model

We use a reduced dynamic mean field (DMF) model (3) to engender lesion effects. The DMF approximates a spiking network model (4, 5) consisting of populations of excitatory and inhibitory neurons with excitatory NMDA synapses and inhibitory GABA synapses. DMF is described by a set of coupled nonlinear stochastic differential equations given below,

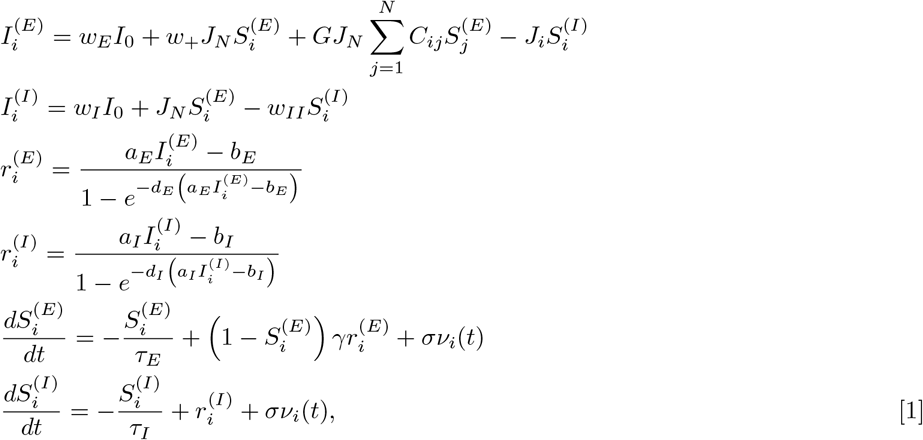

where 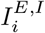 is the input current to area *i* and superscripts represent excitatory (*E*) and inhibitory(*I*) populations in that area. 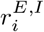 is the population firing rate of excitatory or inhibitory populations of area *i*. 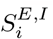 is the average excitatory (inhibitory) synaptic gating variable of area *i. I*_0_ is the effective external input scaled by *w*_*E*_ and *w*_*I*_ for excitatory and inhibitory populations. *w*_+_ is the local excitatory recurrence, *J*_*N*_ is the excitatory synaptic coupling, and *J*_*i*_ is the local feedback inhibitory synaptic coupling. *w*_*II*_ is the local inhibitory recurrence. *C*_*ij*_ is the *i, j*^*th*^ entry in the SC matrix, obtained from diffusion imaging, that scales the long-range excitatory currents between *j*^*th*^ and *i*^*th*^ regions. *G* represents global coupling strength which scales long-range excitatory connections. Descriptions of the parameters and their default values are given in Table S2. To find optimal *G*, the DMF model is simulated for different values of *G*. The optimal value of *G* is chosen based on the highest correlation between empirical and simulated FC, when the excitatory firing rate sustains at *∼*4Hz within all brain regions (6, 7). Stochasticity is incorporated into the two gating variables by additive white Gaussian noise, *σν*_*i*_(*t*), where *σ* is the noise intensity.

**Table S1.** Model parameters and their values.

| Parameter | Description | Value/ Units |
| --- | --- | --- |
| $I_0$ | External input current | 0.382 nA |
| $w_E$ | Weight for excitatory populations | 1 |
| $J_N$ | Long-range excitatory synaptic coupling constant | 0.15nA |
| $w_+$ | Strength of local excitatory recurrent connections | 1.4 |
| $a_E$ | Parameter for input-output function | 310nC<sup>-1</sup> |
| $b_E$ | Parameter for input-output function | 125 Hz |
| $d_E$ | Parameter for input-output function | 0.16s |
| $w_I$ | Weight for inhibitory populations | 0.7 |
| $a_I$ | Parameter for input-output function | 615 nC<sup>-1</sup> |
| $b_I$ | Parameter for input-output function | 177 Hz |
| $d_I$ | Parameter for input-output function | 0.087s |
| $w_{II}$ | Strength of local inhibitory recurrent connections | 1nA |
| $\gamma$ | Learning rate | 0.641 |
| $\tau_E$ | Excitatory time constant | 100 ms |
| $\tau_I$ | Inhibitory time constant | 10 ms |
| $\sigma$ | Noise amplitude | 0.001nA |
| $G$ | Global coupling strength | 0.55 |

### Definitions and descriptions

#### Feedback inhibition control (FIC)

FIC algorithm, proposed by Deco et al. (5), is a recursive process to establish and maintain E-I balance in individual and across all cortical subunits. By the term E-I balance, it means that the average input current 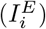 to an excitatory pool of i^*th*^ region is equal to 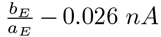, with a tolerance of 0 005. This range of input current clamps the firing rate between 2.63 − 3.55 Hz. When the input current goes beyond the tolerance level, we increase that area’s local feedback inhibitory weight (*J*_*i*_) by a small value (∆). Analytically, when 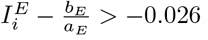, we increase or upregulate the corresponding local feedback inhibition, *J*_*i*_ = *J*_*i*_ + ∆ of the area *i*; otherwise we downregulate the corresponding inhibitory weights *J*_*i*_ = *J*_*i*_ *−* ∆. The process is repeated until all regions’ firing rates reach a critical firing rate regime (2.63 − 3.55 Hz). We simulated the model with the FIC algorithm at different time windows of 10 *sec*.

#### Virtual focal lesion

The virtual focal lesion is introduced into an individual subject’s structural connectome or anatomical topology by the targeted removal of a single node. Specifically, all connections to and from the focal lesioned site have been set to zero in the SC matrix. Thus, a lesioned center is isolated from its neighbors and becomes functionally non-interactive with the remaining intact network. The anatomical topology of the remaining network is kept invariant, except the lesioned node is functionally isolated. The lesioned site will not participate in or influence the functions (or dynamics) of the remaining intact network. A single node is removed to project the exclusive impact of a specific lesioned site. The rest of the remaining network is used in the simulation. Here we consider that the lesion only involves the deletion of a node (‘gray matter’) and its afferent connections. In contrast, we do not attempt to model ‘while-matter’ volume, e.g., including lesions of ‘fibers of passage’ (8). We consider all 68 regions as lesion centers covering the whole cerebral cortex.

#### Virtual lesion model

When the DMF model is put on top of the virtually lesioned SC, we labeled them as the virtual lesion model. Individual node dynamics are governed by the stochastic DMF model spatially coupled via lesioned SC matrix. In principle, any other form of lesioned SC (real, virtual), if incorporated into the dynamical model, can produce similar effects of lesion depending on the lesion type, location, and extent.

**Table S2.** List of all 34 ROIs in each hemisphere. ROI ID represents the order of ROIs in the structural and functional connectivity matrices for each hemisphere.

| Area index | Abbreviation | Full form |
| --- | --- | --- |
| 1 | BSTS | Banks Of Superior Temporal Sulcus |
| 2 | CAC | Caudal Anterior Cingulate |
| 3 | CMF | Caudal Middle Frontal |
| 4 | CUN | Cuneus |
| 5 | ENT | Entorhinal |
| 6 | FITS | Fusiform |
| 7 | IP | Inferior Parietal |
| 8 | IT | Inferior Temporal |
| 9 | ISTH | Isthmus Cingulate |
| 10 | LOCC | Lateral Occipital |
| 11 | LOF | Lateral Orbito Frontal |
| 12 | LING | Lingual |
| 13 | MOF | Medial Orbito Frontal |
| 14 | MT | Middle Temporal |
| 15 | PARH | Parahippocampal |
| 16 | PARC | Paracentral |
| 17 | POPE | Pars Opercularis |
| 18 | PORB | Pars Orbitalis |
| 19 | PTRI | Pars Triangularis |
| 20 | PCAL | Pericalcarine |
| 21 | PCNT | Post Central |
| 22 | PC | Posterior Cingulate |
| 23 | PREC | Precentral |
| 24 | PCUN | Precuneus |
| 25 | RAC | Rostral Anterior Cingulate |
| 26 | RMF | Rostral Middle Frontal |
| 27 | SF | Superior Frontal |
| 28 | SF | Superior Parietal |
| 29 | ST | Superior Temporal |
| 30 | SMAR | Supra Marginal |
| 31 | FP | Frontal Pole |
| 32 | TP | Temporal Pole |
| 33 | TT | Transverse Temporal |
| 34 | INS | Insula |

**Fig. S1.**
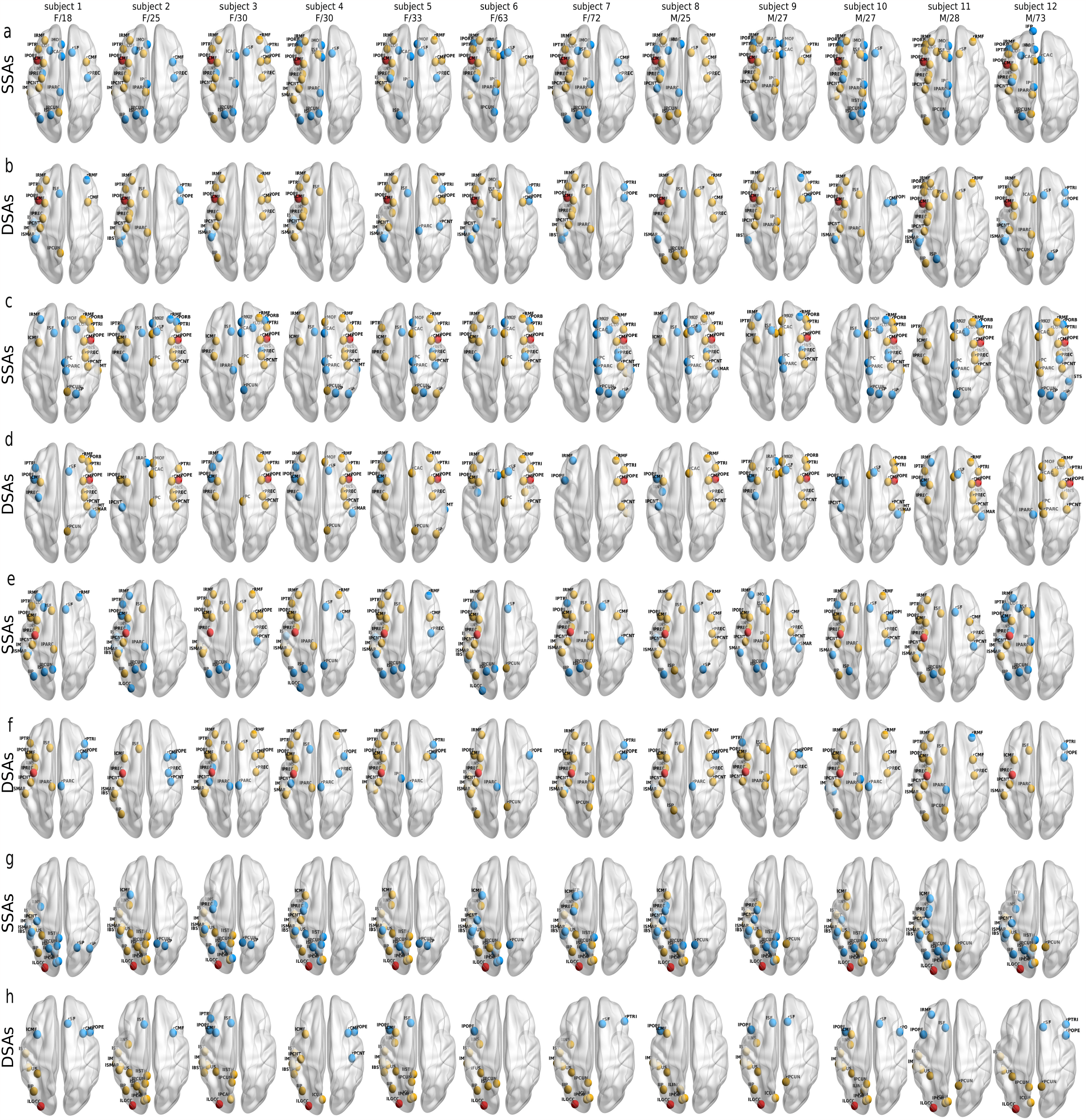
SSAs and DSAs for ten individual subjects corresponding to lesion centers at (a-b) lPOPE, (c-d) rPOPE, (e-f) lPREC, and (g-h) lLOCC. Red sphere represents lesion location. Common areas found in SSAs and DSAs, are shown in yellow, and unmatched areas in blue. Subjects’ gender/age are written in the top of each brain.

**Fig. S2.**
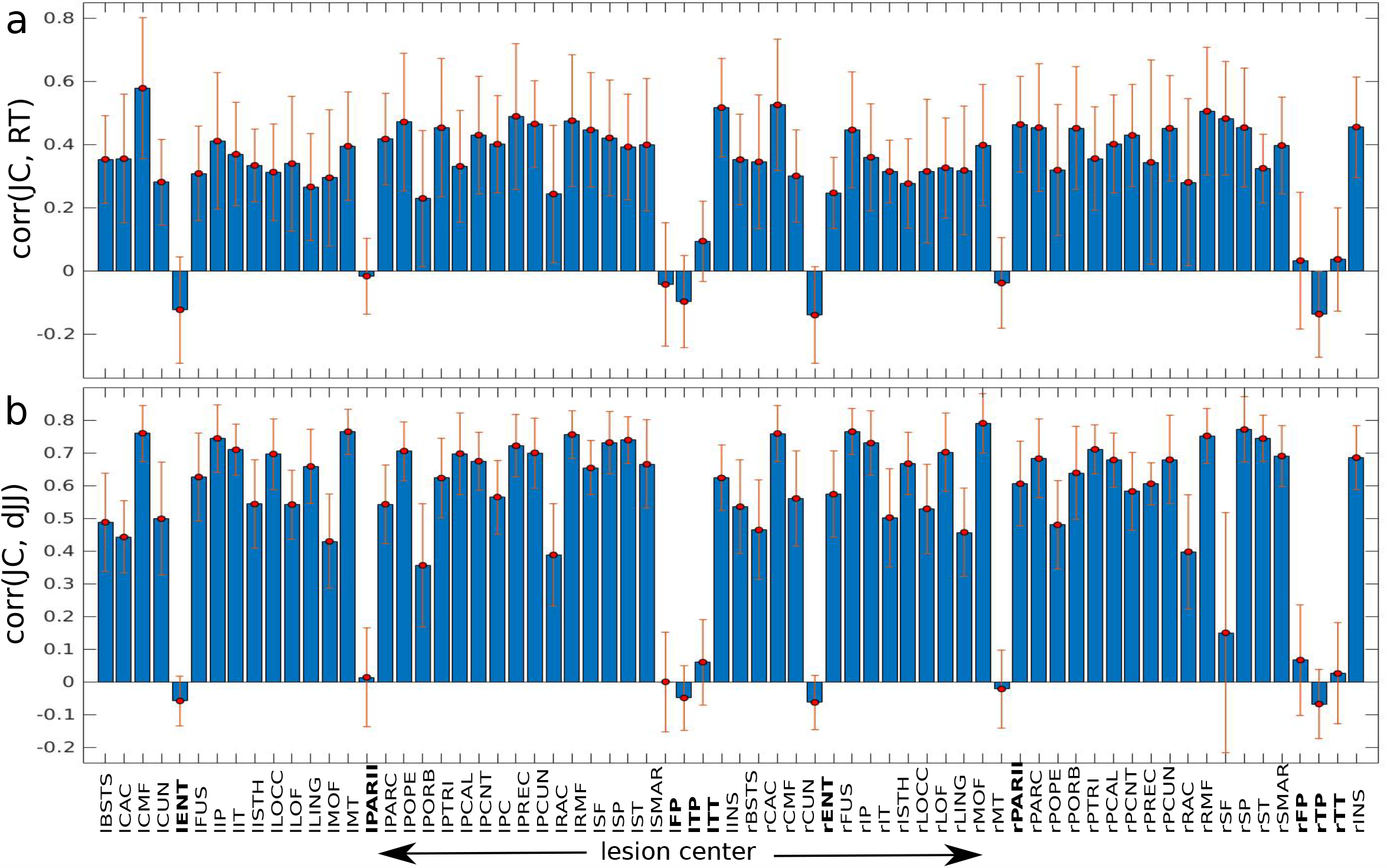
Association between structural and dynamical measures for different lesion centers and subjects. (a) Correlation between *JC* and *RT* has been determined over all the subjects for different lesion centers. The lesion center at a higher degree area displays a positive correlation, except the regions with lower connections show less or negative correlations. Blue bars are obtained by averaging the correlation over the subjects. Subject-wise variations are depicted by red error bars. (b) Similar trends in correlations between *JC* and *dJ J* are observed. Lesion centers are listed below the figure. The areas with low or negative correlations are marked in bold.

### Details of the regions with lower connectivity

We also investigate the anatomical biases of the nodes with low and negative correlations. The distribution of *dJJ* for a single subject at the lesion center, lENT, is shown in SI Appendix, Fig. S3c. The nodes do show very small changes due to lesions at lENT. We also looked at the anatomical connections of the lENT (SI Appendix, Fig. S3d). The brain area lENT is sparsely connected to the rest of the network. We further analyzed all the areas’ average anatomical strength and degree distributions (SI Appendix, Figs. S3e,f).

**Fig. S3.**
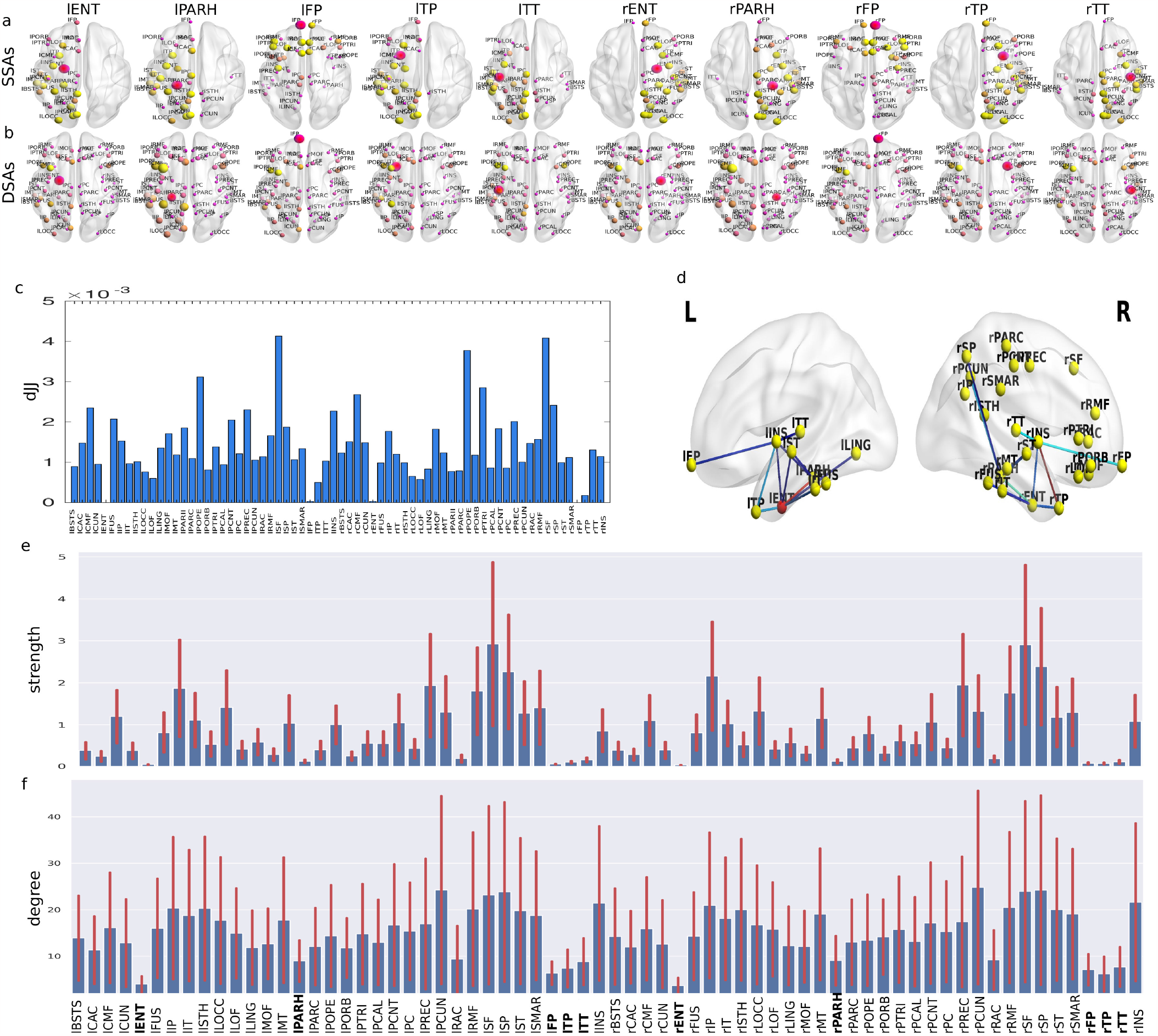
The regions written in bold in Fig. S2 are analyzed at the group level. Findings of SSAs and DSAs for a few lesion sites are given in (a) and (b), respectively. Lesion centers are written on the top of each glass brain plot. Lesion at lENT is an example for a single subject, depicting the distribution of *dJ J* across all areas in (c), and anatomical connections of the lENT in (d). (e) Average anatomical strength and (f) degree of all the areas. Regions with lower connectivities and node weights are marked in bold.

### Group level analysis of SSAs and DSAs

**Fig. S4.**
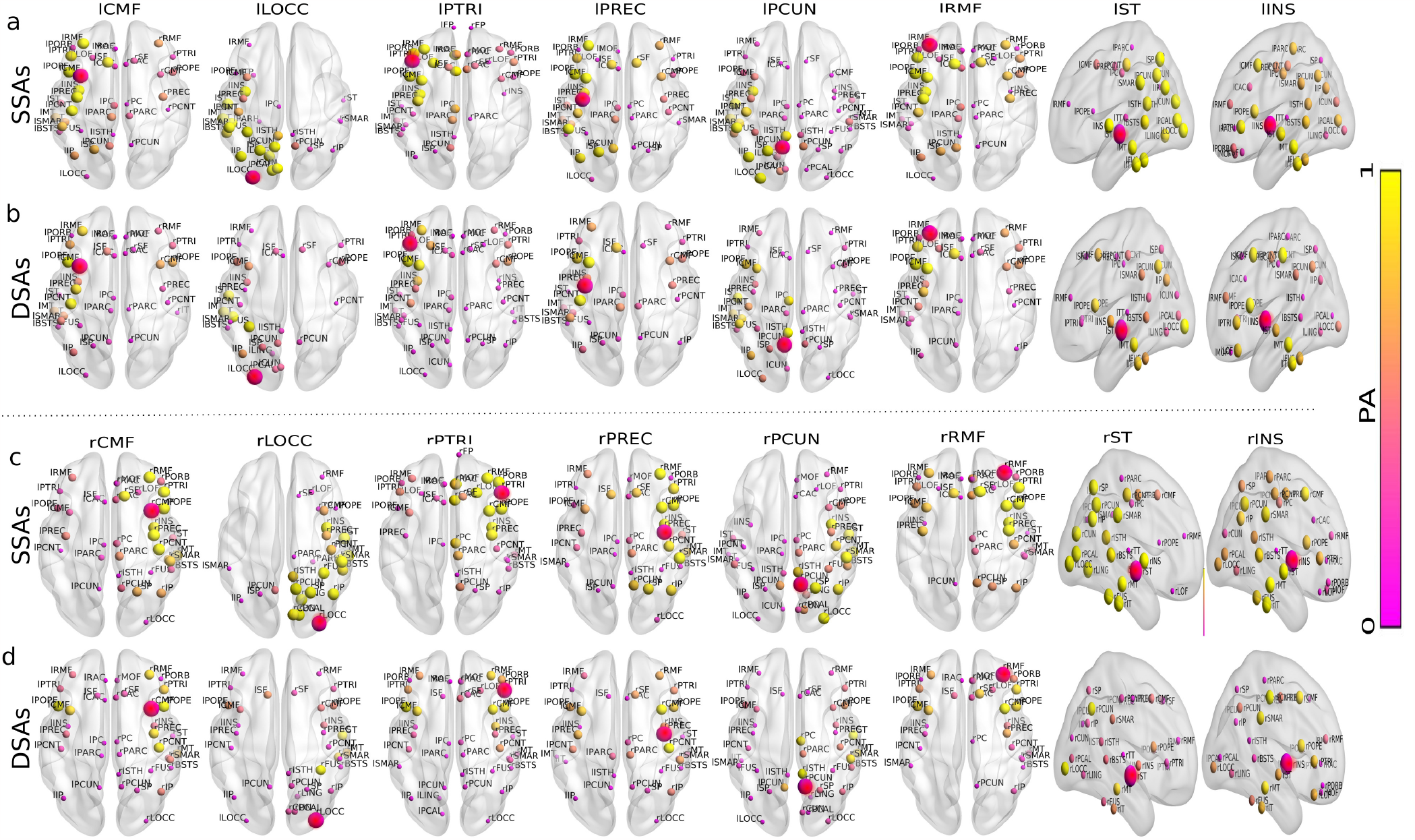
Group level analysis. The identified SSAs and DSAs corresponding to different lesion centers are shown in (a,c) and (b,d), respectively. Abbreviations of lesion sites are written on the top of each brain plot. The red sphere is the lesion site. Yellow areas have higher PA, as they are in almost all subjects. Areas in pink with lower PA indicate that those areas are less probable to be the potential compensatory candidates in all subjects.

### ROI-wise FC analysis over all subjects for different lesion centers

**Fig. S5.**
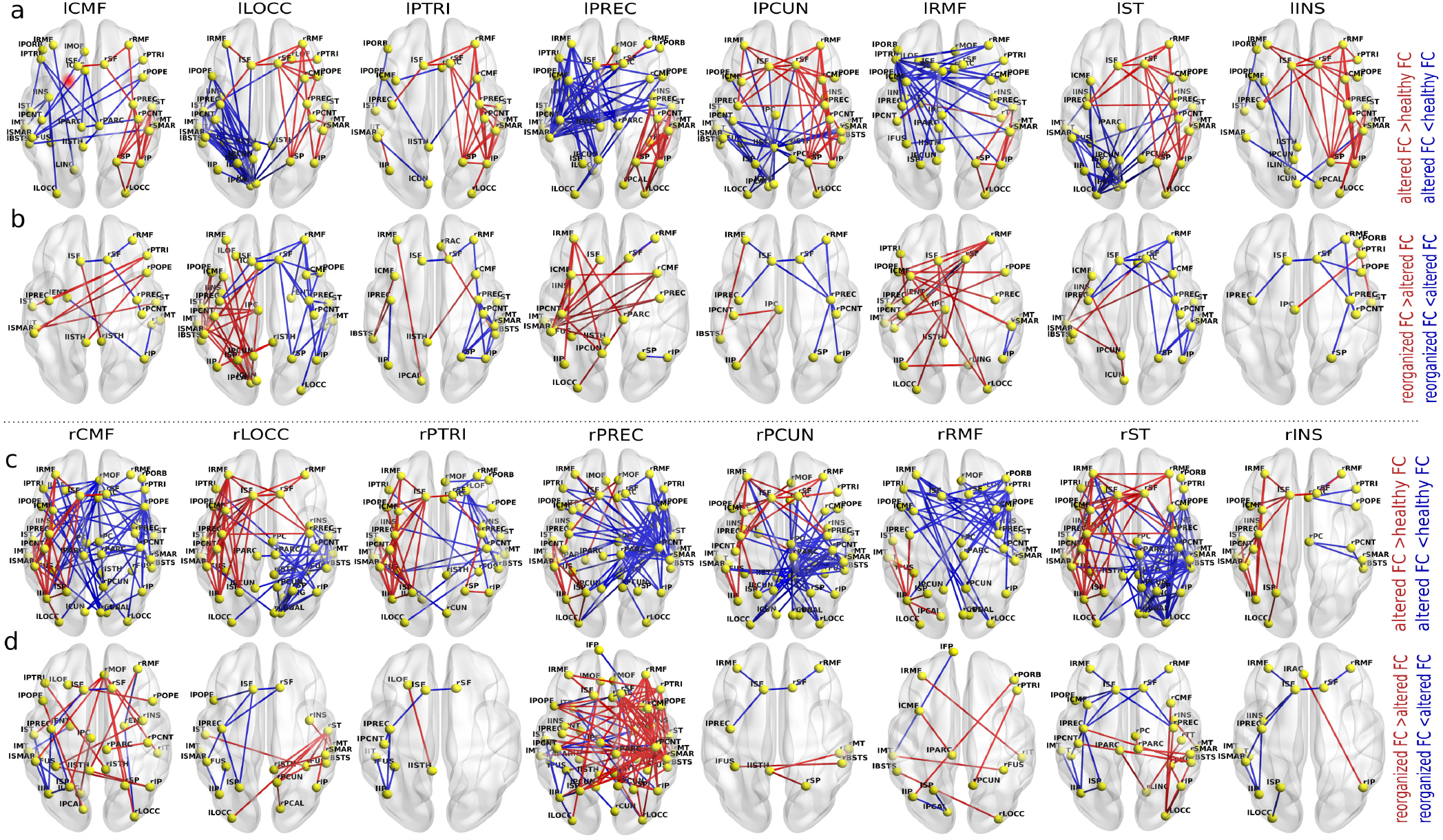
ROI-wise FC analysis for different lesion centers. Significantly changed and rewired links with associated regions are shown in (a,c) and (b,d), respectively. Lesion centers abbreviations are written on the top of each brain. Red and blue edges represent significant increase and decrease in cohesion, respectively.

(16, 17)

**Table S3.**
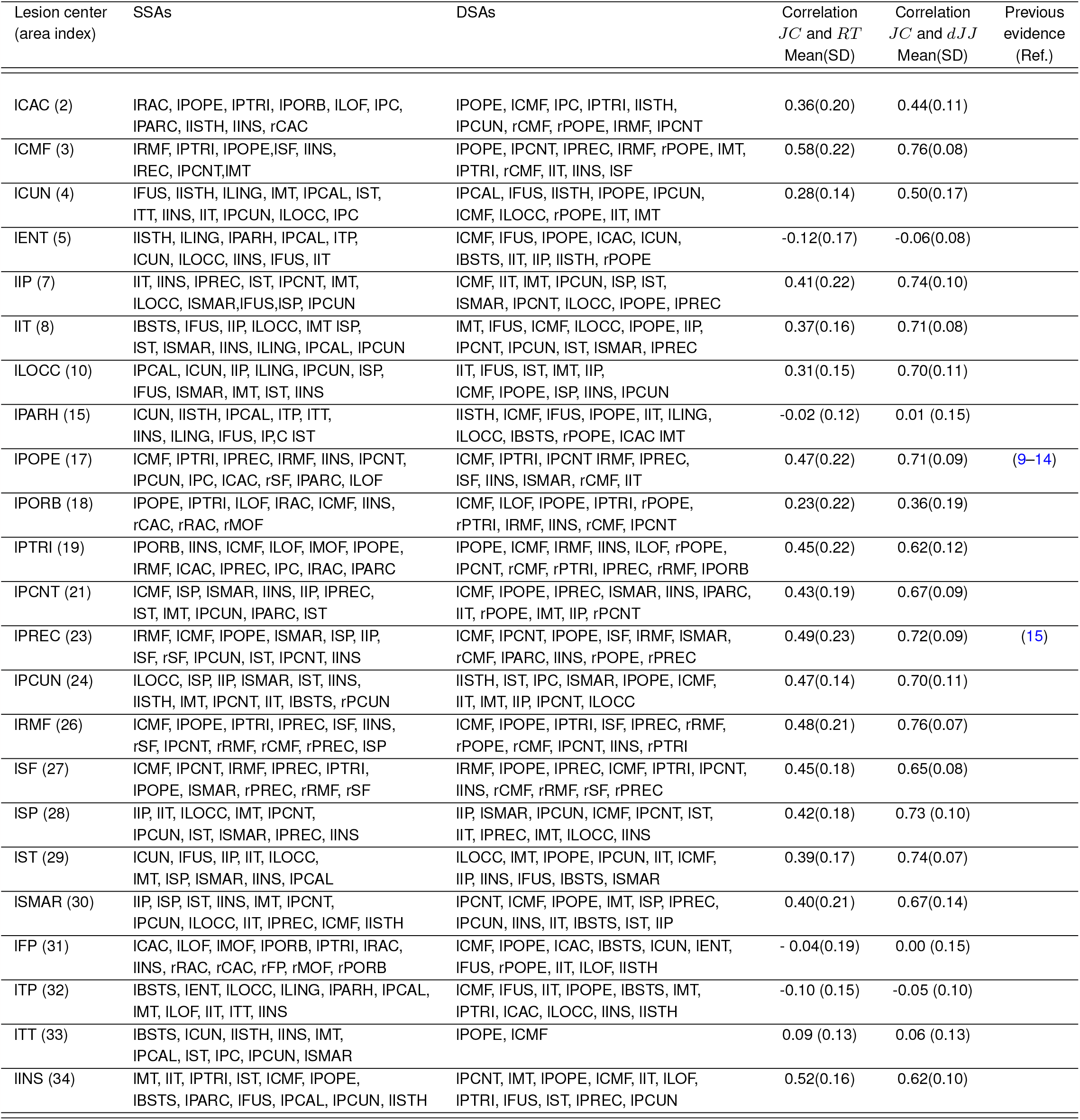
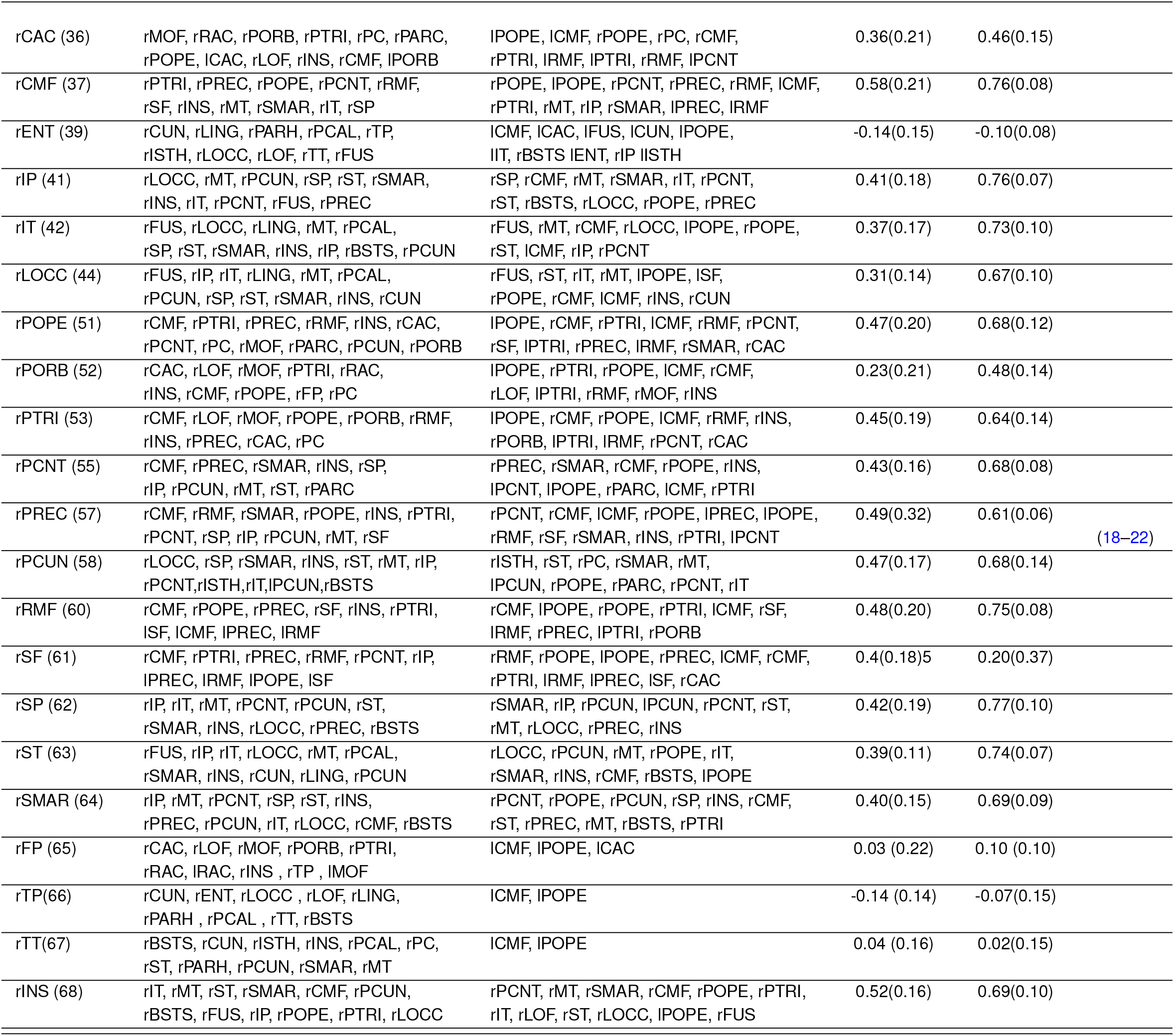
Lesion centers, SSAs, DSAs and correlation between *JC* and *RT*. We choose only those areas as SSAs and DSAs, which have *PA >* 0.8.

### Correlation between JC and other properties of SC

If the nearest neighbours of a node are also directly connected to each other they form a cluster. The clustering coefficient quantifies the number of connections that exist between the nearest neighbours of a node as a proportion of the maximum number of possible connections18

**Fig. S6.**
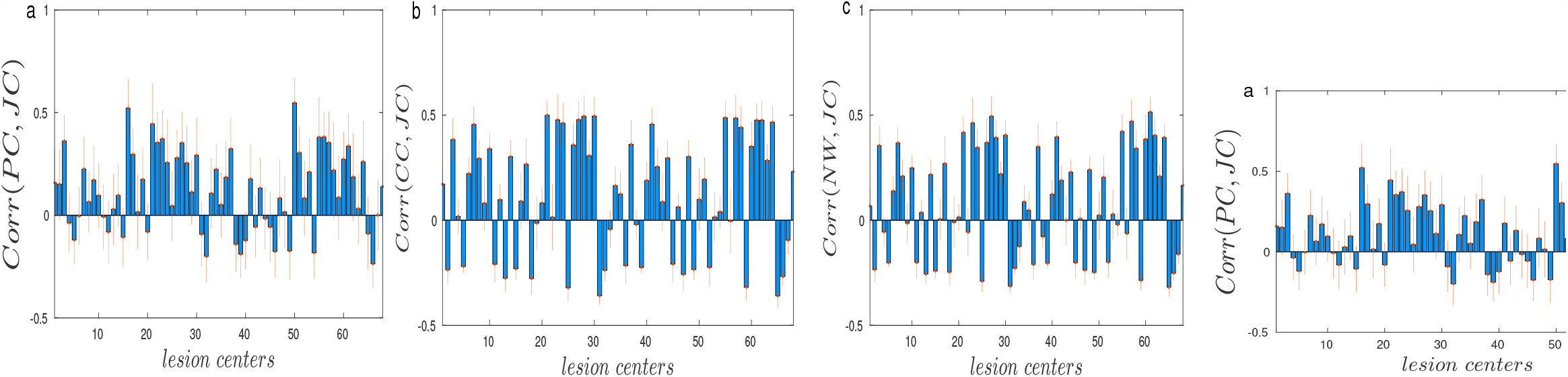
Correlation between JC and other network measures. Mean correlation values are obtained averaging over subjects for different lesion centers. No specific or conclusive patterns are observed in the correlations between JC-PC, JC-CC and JC, NW. PC, CC and NW are participation coefficient, clustering coefficient and node weight. All the network measures are obtained using BCT tools box.

### Functional network properties under three given conditions

The alteration in functional brain network due to structural damage as an immediate impact of lesion and FC re-organization after restoring E-I balance are reconfirmed by measuring several network properties, such as transitivity, average characteristic path length, modularity, and global efficiency, presented in Fig. S7. The network properties computed from healthy, altered, and re-organized FCs are shown in yellow, red, and blue bars, respectively, see Figs. S7(a-d). Transitivity, average characteristic path length, and modularity significantly increased in the altered FCs, when they deviated from the healthy condition due to lost E-I homeostasis. However, the FCs regain nearhealthy conditions when the homeostatic balance is re-established after the lesion. The three network properties are significantly decreased in the re-organized FCs compared to the altered FCs. Conversely, global efficiency significantly decreased (*p <* 0.001) in the altered FC and increased after the re-organization. Alteration/re-organization is further reconfirmed by the probability distributions of FC weights from the three conditions. Distributions for healthy, altered, and re-organized FCs of a single subject are shown in purple, red, and green in Fig. S7e. Blue, red, and green lines indicate the mean values of the three distributions, respectively.

**Fig. S7.**
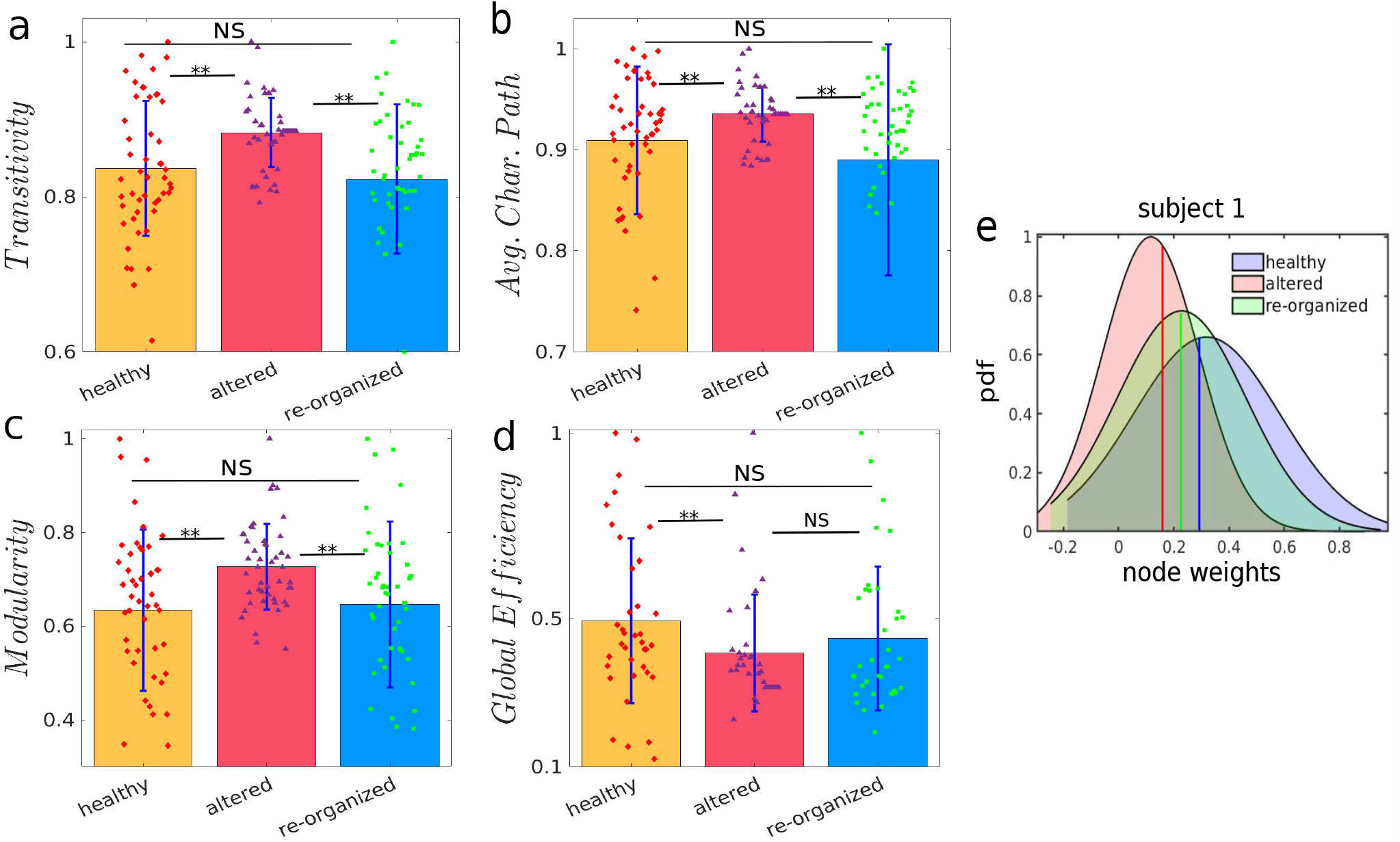
Functional network properties. (a) Transitivity, (b) average characteristic path length, (c) modularity, and (d) global efficiency derived from healthy, altered, and re-organized FCs are plotted in yellow, red, and blue bars, respectively. Deviation over the subjects is shown by the error bar. (e) Probability distributions of the three FCs, driven from the three conditions, confirm FC reshaping after lesion. *∗ ∗ p <* 0.001, ‘NS’ for not significant. Network measures are obtained using BCT tools box.

#### Definition of network properties

i. Transitivity (23), or clustering coefficient, measures the tendency of the nodes to cluster together. High transitivity means that the network contains communities or groups of nodes that are densely connected internally. Transitivity of a graph with degree sequence *k* is 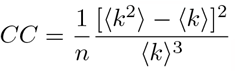, where ⟨*k*⟩ = 1/*n* Σ_*i*_ *k*_*i*_ is the mean degree and 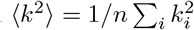 is the mean square degree.
ii. Characteristic path length (24), 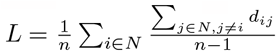, where *N* is the set of all nodes; *n* is the total nodes; *d*_*ij*_ is the weighted shortest path length between *i* and *j*. The characteristic path length for weighted graphs is an estimate of proximity. The global efficiency is the average of the inverse shortest path length and is inversely related to the characteristic path length. The local efficiency is the global efficiency computed on the node’s neighborhood and is related to the clustering coefficient.
iii. Modularity (25), 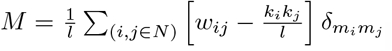, where *l* is the sum of weights in the network; *w*_*ij*_ is the connectivity weight between nodes *i* and *j*; *k*_*i*_, *k*_*j*_ are the weighted degrees of nodes *i* and *j*, respectively. Modularity gives network resilience and adaptability, measuring the degree of segregation. Communities are subgroups of densely interconnected nodes sparsely connected with the rest of the network. In the case of a functional network, modularity signifies coherent clusters of functional modules.
iv. Global efficiency (26), 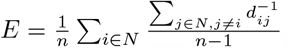, where *N* is the set of all nodes; n is the total number of nodes; *d*_*ij*_ is the weighted shortest path length between node *i* and *j*. It captures the integration property of a network.

## Notes

### Competing Interest Statement

The authors have declared no competing interest.

### Summary of Updates

The version of the manuscript is revised to update the figures and paragraphs under a few sections.

